# Signal peptide mimicry primes Sec61 for client-selective inhibition

**DOI:** 10.1101/2022.07.03.498529

**Authors:** Shahid Rehan, Dale Tranter, Phillip P. Sharp, Eric Lowe, Janet L. Anderl, Tony Muchamuel, Vahid Abrishami, Suvi Kuivanen, Nicole Wenzell, Andy Jennings, Chakrapani Kalyanaraman, Gregory B. Craven, Tomas Strandin, Matti Javanainen, Olli Vapalahti, Matt Jacobson, Dustin McMinn, Christopher J. Kirk, Juha T. Huiskonen, Jack Taunton, Ville O. Paavilainen

## Abstract

Preventing the biogenesis of disease-relevant proteins is an attractive therapeutic strategy, but attempts to target essential protein biogenesis factors have been hampered by excessive toxicity. Here, we describe KZR-8445, a cyclic depsipeptide that targets the Sec61 translocon and selectively disrupts secretory and membrane protein biogenesis in a signal peptide-dependent manner. KZR-8445 potently inhibits the secretion of proinflammatory cytokines in primary immune cells and is highly efficacious in a mouse model of rheumatoid arthritis. A cryo-EM structure reveals that KZR-8445 occupies the fully opened Se61 lateral gate and blocks access to the lumenal plug domain. KZR-8445 binding stabilizes the lateral gate helices in a manner that traps select signal peptides in the Sec61 channel and prevents their movement into the lipid bilayer. Our results establish a framework for the structure-guided discovery of novel therapeutics that selectively modulate Sec61-mediated protein biogenesis.

## INTRODUCTION

Secretory and integral membrane protein biogenesis occurs primarily at the surface of the endoplasmic reticulum (ER). Such proteins, including cytokines and cell surface receptors, comprise approximately one quarter of the eukaryotic proteome^1^ and play critical roles in cancer and inflammatory diseases. Biogenesis of most membrane and secreted proteins requires the Sec61 translocon^2^, an evolutionarily conserved transmembrane channel complex that facilitates translocation and membrane integration of nascent polypeptides. As a key nexus of the secretory pathway, pharmacological modulation of Sec61 could potentially prevent the biogenesis of proteins, with critical roles in intercellular communication and disease physiology.

A defining feature of Sec61 clients is the presence of an N-terminal signal peptide or transmembrane helix which is required for cotranslational ER targeting and insertion^3^. Signal peptides (typically 15-45 amino acids in length) lack a clear consensus motif but generally consist of an N-terminal region with polar residues, a central hydrophobic region that binds transiently to Sec61, and a C-terminal region containing the signal peptidase cleavage site^4^. A nascent protein’s signal peptide or N-terminal transmembrane segment exits the ribosome and initially engages the cytosolic signal recognition particle (SRP)^5^. This ribosome-nascent chain complex (RNC) is then targeted to the ER membrane and delivered to the Sec61 translocon. RNC docking has been proposed to alter the conformation of Sec61 to prime it for polypeptide insertion^2^. A structure of mammalian Sec61 in complex with an inserted signal peptide^6^ revealed that the hydrophobic region intercalates between Sec61 transmembrane helices (TM) (the “lateral gate”) and thereby gains access to the lipid bilayer. Hence, a primary mechanism by which signal peptides promote opening of the Sec61 channel is through dynamic hydrophobic interactions with residues of the lateral gate helices TM2 and TM7. Despite the stringent requirement for recognition by the Sec61 complex, signal peptides are remarkably diverse in sequence and therefore may interact with Sec61 lateral gate helices in different ways during the dynamic translocation process.

Consistent with its essential cellular role, complete blockade of Sec61-mediated protein import with small-molecule inhibitors is eventually toxic to mammalian cells^7–11^. A few Sec61 inhibitors, including mycolactone, apratoxin and their derivatives have been tested in preclinical disease models (primarily cancer). However, these compounds lack Sec61 client selectivity and have generally suffered from a low therapeutic index, precluding further development^12^. By contrast, certain cotransin cyclic heptadepsipeptides potently inhibit Sec61 in a client-selective manner^13,14^. The main determinant of client-selective inhibition appears to be the primary amino acid sequence and corresponding biophysical properties of the ER-targeting signal peptide or the N-terminal transmembrane segment. Previous work demonstrated that cotransin directly targets the Sec61α subunit of the heterotrimeric Sec61 channel^15^, and unbiased mutagenesis screens suggested that cotransins bind Sec61α near the lumenal plug and the lateral gate^16,17^. Remarkably, modifications to the cotransin structure can alter the range of inhibited Sec61 clients^18^, suggesting the possibility of identifying cotransin variants that affect distinct subsets of secretory and membrane proteins. However, the lack of a molecular-level understanding of how cotransins bind Sec61 to achieve substrate-selective effects has prevented the rational design of cotransins with altered pharmacological profiles.

Here, we present a cryogenic electron microscopy (cryo-EM) structure of Sec61 bound to KZR-8445, a novel cotransin variant that prevents biogenesis of a subset of Sec61 clients and is efficacious in a mouse model of rheumatoid arthritis. Our cryo-EM structure reveals that KZR-8445 binds to the central region of the Sec61 lateral gate in an open configuration, with direct contacts to the lumenal plug. Similar to nascent signal peptides, KZR-8445 disrupts the interhelical lateral gate interactions that stabilize the channel in its closed configuration. Based on our structure, we propose a mechanism in which inhibitor binding to the open lateral gate and closed lumenal plug specifically traps nascent signal peptides in the cytosolic vestibule, thus providing a framework for the design of new cotransins with increased levels of client discrimination.

## RESULTS

### KZR-8445 is a client-selective Sec61 inhibitor with therapeutic efficacy in vivo

The previously identified cotransins, CT8 and PS3061, potently block the secretion of pro-inflammatory cytokines^18^ and inhibit viral replication^19,20^, including SARS CoV-2^21^. To test whether cotransins can serve as disease-modifying agents in vivo, we designed KZR-8445 (**Fig. 1A**), a fluorinated variant of PS3061 with improved physicochemical properties and pharmacokinetics. Similar to PS3061, KZR-8445 blocked CoV-2 replication and virus-induced cytotoxicity in Vero cells, an effect that was likely mediated by inhibition of spike protein biogenesis (**Supplementary Fig. 1A–E**). KZR-8445 was also recently shown to overcome dexamethasone resistance in T-cell acute lymphoblastic leukemia cells in vitro^22^.

**Figure 1.**
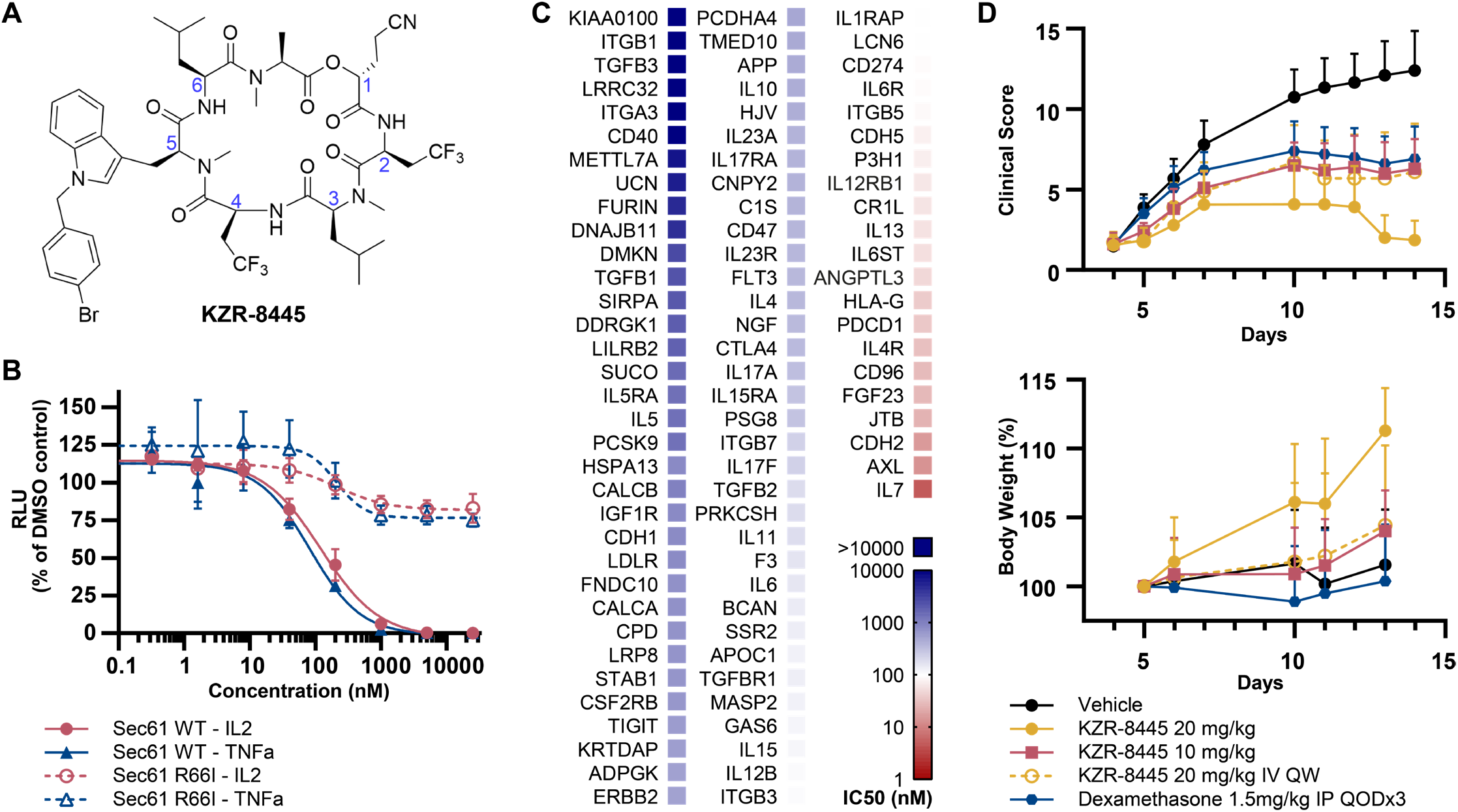
KZR-8445 is a client-selective Sec61 inhibitor and is efficacious in a mouse arthritis model. **(A)** Chemical structure of KZR-8445. **(B)** Cells stably expressing WT or R66I Sec61α were transfected with dox-inducible Gaussia luciferase (GLuc) reporter constructs fused to the C-terminus of IL2 or TNF. After treatment with doxycycline and the indicated concentrations of KZR-8445 for 24 h, GLuc activity was quantified. Data are depicted as mean from three biological replicates ± s.d. **(C)** Cells were transfected with dox-inducible GLuc constructs fused to the indicated human signal peptides (SP-GLuc), treated with doxycycline and increasing concentrations of KZR-8445 for 24 h, and assessed for GLuc activity. The heat map depicts IC50 values calculated from each SP-GLuc dose-response curve. (n=3 for IC50 determination) **(D)** Arthritis was induced in BALB/c mice with antibodies specific for type II collagen (mAb) on day 0 and endotoxin on day 3. On day 4, when disease was present in all animals, mice were randomized and treated for 2 weeks with thrice weekly (QODx3) or weekly (QW) intravenous administration of vehicle or KZR-8445 or intraperitoneal administration of dexamethasone. Clinical scores (0-4 per paw; n = 10/group) and body weights were followed until day 14. Data are presented as mean values ± SEM.

To investigate the effects of KZR-8445 on secretory and membrane protein biogenesis, and to establish Sec61α as the relevant target, we quantified secretion of a doxy-cycline-inducible Gaussia luciferase (GLuc) reporter fused to the C-terminus of two full-length pro-inflammatory cytokines, IL2 (which contains a cleavable signal peptide) and TNFα (which contains an N-terminal transmembrane signal anchor). We performed the GLuc secretion assays in cells stably expressing exogenous wild type (WT) Sec61α or a plug domain mutant (R66I) which was previously shown to confer resistance to CT8 and a cotransin natural product^16,17^ (**Fig. 1B**). KZR-8445 potently blocked secretion of both GLuc reporter constructs (IC50 ∼100 nM), and this effect was reduced in cells expressing R66I Sec61α. Hence, KZR-8445 acts directly on Sec61α to block protein secretion. KZR-8445 is structurally related to CT8, which discriminates among Sec61 clients based largely on their N-terminal signal peptide18. To test whether KZR-8445 exhibits signal peptide-dependent selectivity, we screened a panel of 89 signal peptide-GLuc reporter constructs derived from diverse disease-relevant Sec61 clients. Results from this screen revealed a striking variation of signal peptide sensitivity to KZR-8445, with IC50s ranging from 5 nM to >25 μM (median 366 nM), depending on the signal peptide (**Fig. 1C**). We also directly compared KZR-8445 to mycolactone A/B, a structurally distinct and relatively client-nonselective Sec61 inhibitor^8,23^, for their ability to discriminate between VCAM and preprolactin (pPL) signal peptides. Whereas KZR-8445 inhibited VCAM-GLuc 20-fold more potently than pPL-GLuc, mycolactone A/B was equipotent against both reporters (**Supplementary Fig. 1F**). These results further highlight a unique signal peptide inhibition profile and suggest that KZR-8445 might exert a distinct mechanism of Sec61 modulation as compared with mycolactone A/B.

Finally, we tested the effects of KZR-8445 on the production of endogenous pro-inflammatory cytokines by activated human peripheral blood mononuclear cells and mouse splenocytes (**Supplementary Fig. 1G, H**). KZR-8445 blocked secretion of 6 out of 7 tested cytokines with submicromolar IC50s, while having minimal effects on cell viability at concentrations up to 20 μM. Notably, secretion of IL1β, which is not Sec61-dependent, was unaffected by KZR-8445.

Although the nonselective Sec61 inhibitor mycolactone A/B showed modest efficacy in a mouse skin inflammation model when tested at its maximum tolerated dose^24^, client-selective Sec61 modulators have not been evaluated in animal models of chronic inflammatory disease to the best of our knowledge. To determine if the anti-cytokine secretion effects observed in vitro would translate to an anti-inflammatory effect in vivo, we tested KZR-8445 in a collagen antibody-induced mouse model of rheumatoid arthritis. When administered thrice weekly after the onset of disease, KZR-8445 blocked disease progression in a dose-dependent manner (10 mg/kg, p=0.01; 20 mg/kg, p<0.0001, one-way ANOVA vs. vehicle). At the highest dose KZR-8445 produced a stronger anti-inflammatory effect than the corticosteroid dexamethasone. Weekly administration of KZR-8445 was also found to be effective (20 mg/kg, p=0.007). Finally, none of the KZR-8445 treatment groups experienced significant toxicity as measured by body weight changes (**Fig. 1D**). Collectively, our data show that KZR-8445 is a potent and substrate-selective inhibitor of Sec61. Unlike other previously described Sec61 inhibitors, KZR-8445 is efficacious and well tolerated in an animal model of rheumatoid arthritis.

### Isolation and cryo-electron microscopy of the ribosome–Sec61 complex bound to KZR-8445

To understand the molecular basis of Sec61 inhibition by KZR-8445, we pursued structural investigations using single-particle cryogenic electron microscopy (cryo-EM). Our initial attempts to isolate cotransin-bound ribosome– Sec61 complexes using the commonly used detergent digitonin were not successful. Several other detergents were screened for retention of native-like cotransin binding by subjecting solubilized Sec61 to a cotransin photoaffinity labeling assay^15^ (**Supplementary Fig. 2C**). This screen identified lauryl maltoside neopentyl glycol (LMNG), a lipid-mimetic detergent that permits maximal photo-cotransin binding to Sec61 as indicated by specific crosslinking to the Sec61α subunit in ER microsomes or LMNG-solubilized ER membranes. Importantly, photo-cotransin crosslinking was efficiently competed not only by excess KZR-8445, but also by a structurally unrelated Sec61 inhibitor apratoxin A^11^, suggesting that LMNG solubilization may allow preparation of Sec61 complexes bound to other inhibitors.

Having established solubilization conditions that preserve native-like KZR-8445 binding to Sec61, we proceeded with isolation of the ribosome/Sec61/KZR-8445 complex for single particle cryo-EM structure determination. We solubilized sheep ER microsomes that were pretreated with 10 μM KZR-8445, isolated the ER-bound 80S polysomes, converted them to 80S monosomes by RNAse treatment, and subjected them to gravity-flow size exclusion chromatography (**Supplementary Fig. 2.1A**). The fractions containing intact ribosome/Sec61 complexes (**Supplementary Fig. 2.1B**) were deposited on holey cryoEM grids with a thin carbon support. We collected a dataset consisting of 30266 micrographs (**Supplementary Table 1**) and carried out single-particle structure determination (**Supplementary Fig. 2.2**). This resulted in a consensus reconstruction with an overall resolution of 3.2 Å and local resolution in the Sec61 region of 2.6–7.0 Å, with poorest density in the N-terminal half of the channel (**Supplementary Fig. 2.3**).

Consistent with previous structural work^6,25,26^, the obtained cryo-EM map revealed a well-defined ribosome density, with additional density representing the trimeric Sec61 translocon proximal to the ribosome exit tunnel (**Fig. 2A, Supplementary Fig. 2.4A**,**B**). The quality of the cryoEM density encompassing the C-terminal half of Sec61α allowed unambiguous modeling within this region, while the weaker density representing the N-terminal half likely reflects the greater conformational flexibility of this part of the channel. In addition to density corresponding to Sec61 subunits, we observed a prominent density in the center of the channel, near the lumenal plug and connected to Sec61 TM7 (**Fig. 2B, Supplementary Fig. 2.4C**). The additional density is proximal to Sec61α residues that, when mutated, were previously found to confer resistance to various cotransin analogs^16,17^ (**Supplementary Fig 2.4D**). The overall shape and size of the presumed KZR-8445 density fits the expected conformation of KZR-8445 as determined by molecular modeling (**Fig. 2C**). Further, we observe the same density in both translating and non translating ribosomes as assessed by the presence of A/P and P/E-site tRNAs, suggesting that the additional density does not belong to a nascent polypeptide. We conclude that the extra density in the Sec61 channel likely represents bound KZR-8445.

**Figure 2.**
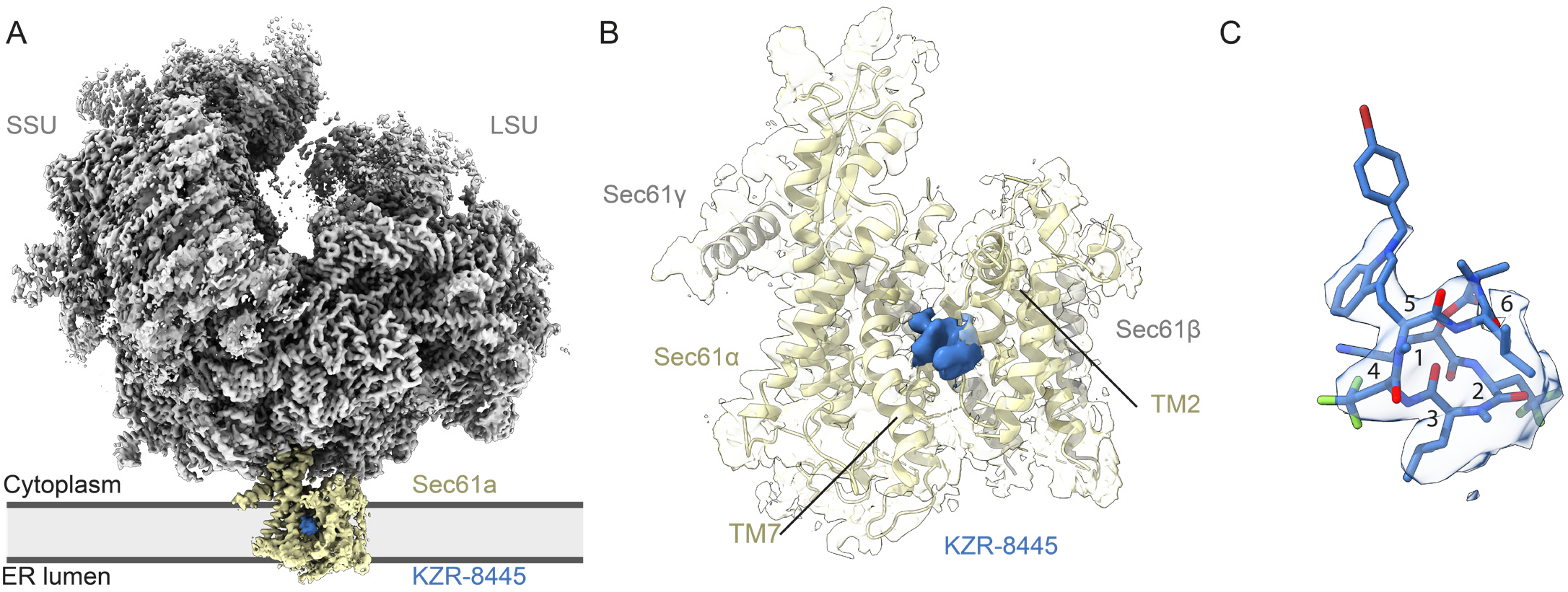
Structure of the mammalian Sec61 translocon with KZR-8445, a substrate-selective translocation inhibitor. **(A)** Cryo-EM map of the mammalian ribo-some-bound heterotrimeric Sec61 translocon in complex with the cotransin analog KZR-8445. The map was low-pass filtered to 4 Å with density features corresponding to the ribosome (LSU and SSU), Sec61, and KZR-8445. **(B)** Additional density assigned to KZR-8445 is located between TM2,TM3, TM7, and lumenal plug helices. KZR-8445 is bound to the center of the lateral gate, which adopts an open conformation. **(C)** Fit of KZR-8445 to the observed density.

### KZR-8445 interacts with the Sec61 lateral gate and the plug domain

To create a model of Sec61 bound to KZR-8445, we used the closed model of mammalian Sec61^25^ as a starting template. Real space refinement and manual modeling revealed that the KZR-8445 macrocycle is inserted into a fully opened lateral gate (**Fig. 3A**). We note that the conformation of the Sec61 lateral gate in our modeled structure is generally similar to that reported for the yeast post-translational translocon^27,28^, as well as the recently described structure of mycolactone A/B bound to a distinct site near the cytosolic vestibule of mammalian Sec61^29^ (**Fig. 4B, Supplementary Fig. 4D**). Therefore the Sec61 conformation we observe represents an energetically favourable state, which may exist during the dynamic protein translocation process.

**Figure 3.**
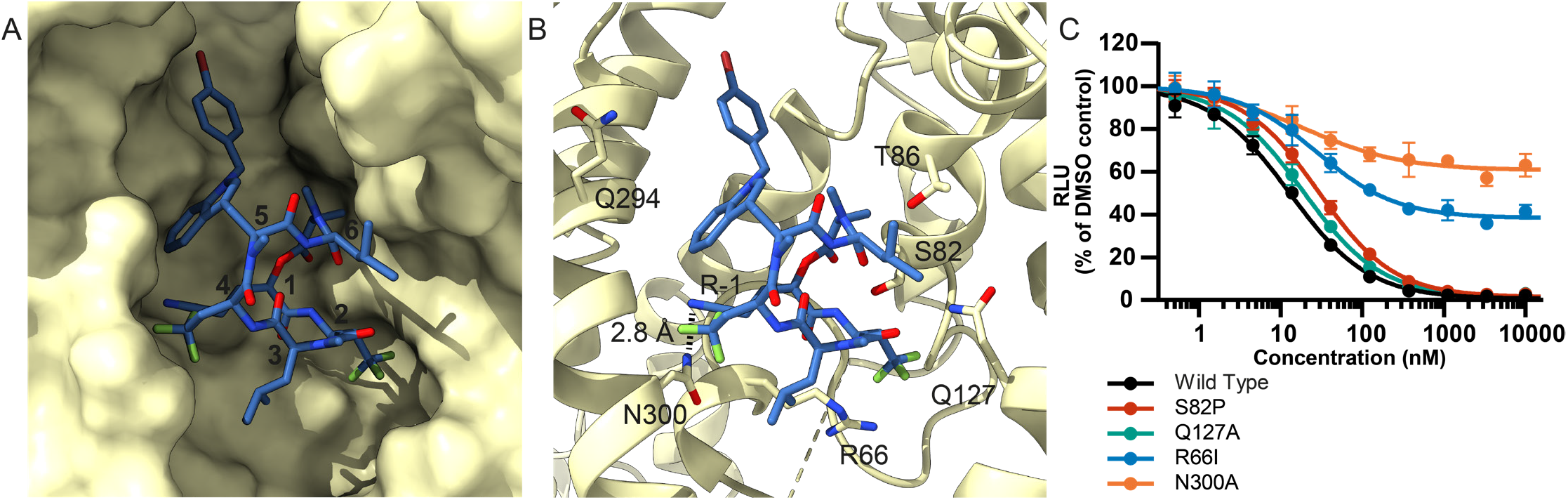
Detailed view of the KZR-8445 binding site. **(A)** Solvent-excluded surface view of the open Sec61α lateral gate bound to KZR-8445. **(B)** Polar residues of the Sec61α cavity proximal to KZR-8445 **(C)** KZR-8445 sensitivity of a VCAM-SP Gluc reporter construct in cells expressing the indicated Sec61α mutant. Data are depicted as mean from four biological replicates ± s.d.

**Figure 4.**
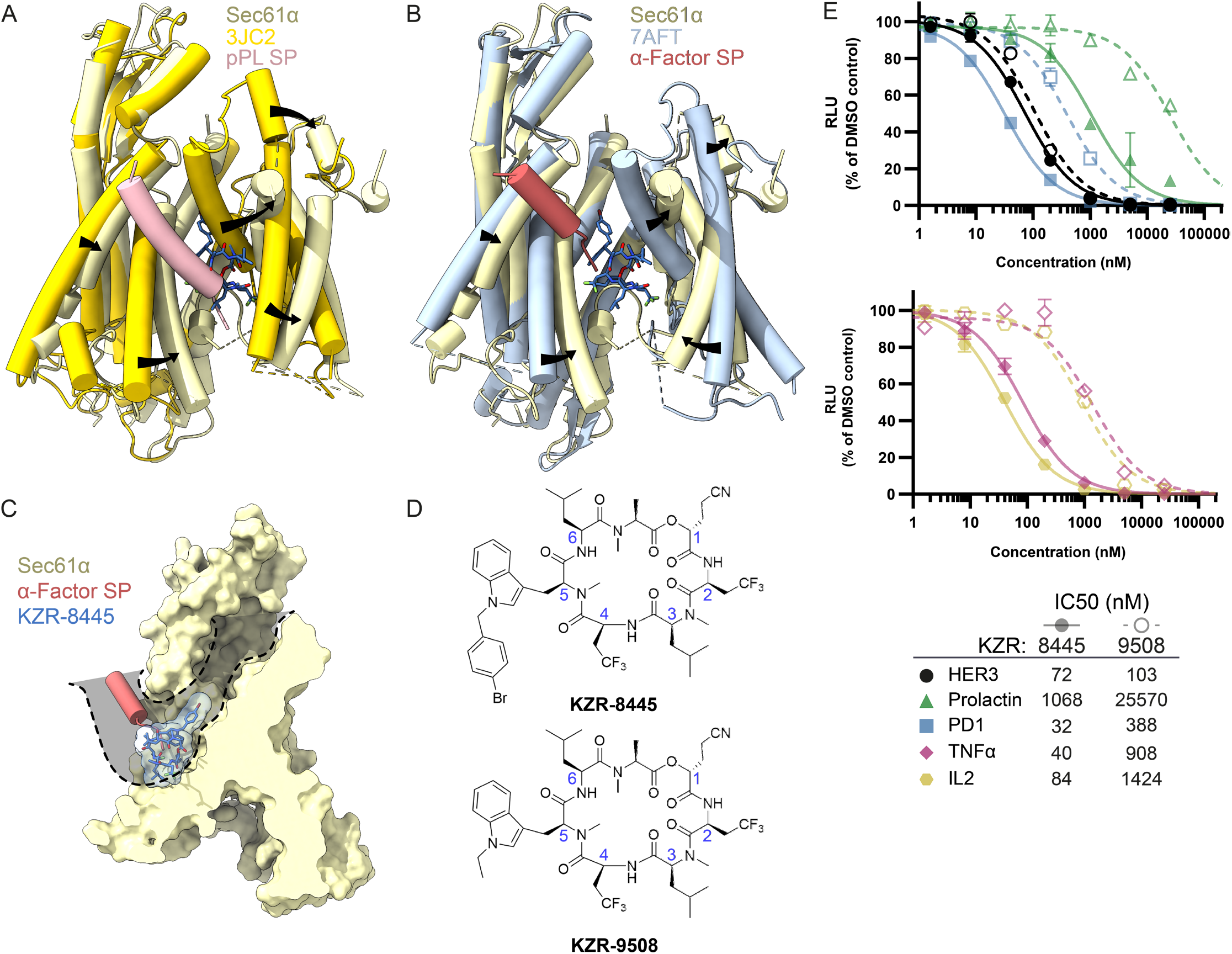
Structural insights lead to improved signal peptide selectivity. **(A and B)** Superimposition of Sec61/KZR-8445 complex with **(A)** preprolactin signal peptide (salmon) engaged with Sec61 (PDB: 3JC2, bright yellow) or **(B)** yeast α-factor signal peptide (red) engaged with Sec61 (PDB: 7AFT, blue). **(C)** Sec61/KZR-8445 superimposed with yeast α-factor signal peptide showing the solvent-excluded surface of the translocon. Indicated in gray is a putative route traversed by nascent signal peptides to reach the binding site occupied by the yeast α-factor signal peptide. **(D)** Structures of KZR-8445 and KZR-9508, a cotransin with a truncated R-5 side chain. **(E)** Cells were stably transfected with dox-inducible Gaussia luciferase (GLuc) reporter constructs fused to the C-terminus of the indicated signal peptides (top), or full-length IL2 or TNFα (bottom). Following treatment with doxycycline and the indicated concentrations of KZR-8445 for 24 h, GLuc activity was quantified. Data are depicted as mean from three biological replicates ± s.d.

We initially modeled KZR-8445 into the central cavity of Sec61 by identifying the likely low-energy conformations of the macrocycle^30,31^. This analysis revealed two distinct backbone conformations of KZR-8445 with the main difference being a *cis* or *trans N*-methyl amide bond linking R-2 and R-3 residues. We fitted both models into the cryo-EM density and obtained a better match with the cis conformation of KZR-8445, which is similar to the conformation of the related cotransin natural product, HUN-7293, as determined by NMR and x-ray crystallography^32^. We therefore used this conformation to model KZR-8445 into the cryo-EM density. Orientation of KZR-8445 was facilitated by assigning a prominent feature of the density to the R-5 bromobenzyl-tryptophan side chain (**Fig. 2C, Supplementary Fig. 2.4C**).

In our structure, KZR-8445 is located in the center of the channel, bounded by Sec61 lateral gate and lumenal plug helices, and the hydrophobic interior of the lipid bilayer (**Fig. 3A and Supplemental Fig 3.1C**). In this arrangement, the channel pore is occluded and access to the lipid bilayer is restricted. The lumenal plug is visible in the cryo-EM density indicating that it is ordered when bound to KZR-8445. We note that the resolution of the KZR-8445 density itself is limited to 4–5 Å and the smaller KZR-8445 side chains are therefore unresolved in our map. However, we consistently observe a salient density that connects KZR-8445 with Sec61α, which we have modeled as a hydrogen bond between the side-chain amide of N300 and the R-1 nitrile of KZR-8445 (**Fig. 3B**). Other proximal Sec61α residues that could potentially form polar interactions with KZR-8445 include S82, T86, and Q127. We note that T86, Q127, and N300 have been suggested to form a polar cluster whose intramolecular interactions maintain the closed state of the channel^33^. Competition with these interactions may contribute to the mechanism by which KZR-8445 opens the Sec61 lateral gate. Consistent with a critical role for N300, ectopic expression of N300A Sec61α in HEK293 cells conferred resistance to KZR-8445 in the secreted GLuc assay (**Fig. 3C and Supplemental Fig. 3.1D**). Although R66 does not form polar contacts with KZR-8445 based on our model, it makes several hydrophobic contacts and mutation of this plug helix residue (R66I) also conferred resistance. By contrast, mutation of other proximal polar residues had little or no effect in this assay. We conclude that Sec61α N300 is essential for KZR-8445-mediated inhibition of protein secretion. N300 may form a hydrogen bond with the R-1 cyanoethyl side chain, although it is also possible that N300 mutations intrinsically favor signal peptide insertion by disrupting interactions between lateral gate helices. Competition with intramolecular interactions may explain how KZR-8445 promotes channel opening in the absence of a signal peptide.

To assess the stability of the proposed KZR-8445 binding mode and to gain additional insights into interactions between KZR-8445 and Sec61α, we used atomistic molecular dynamics (MD) simulations. Two complementary sets of simulations using different molecular force fields suggest that the modeled pose of KZR-8445 is stable as it does not move out of the modeled binding pocket during the simulation (**Supplementary Fig. 3.2 A to C**). The presence of KZR-8445 between TM2 and TM7 maintains the lateral gate in an open conformation, whereas removal of KZR-8445 leads to its closure within hundreds of nanoseconds (**Supplementary Fig. 3.2C and 3.2D**). While KZR-8445 binding to Sec61 is mainly governed by van der Waals interactions, our analysis also suggests the formation of hydrogen bonds with Sec61α residues N300 and Q127. Finally, the overall Sec61 structure shows similar root mean square displacement values both with and without KZR-8445 in our microsecond-long simulations, indicating that the inhibitor has little effect on Sec61 stability apart from the lateral gate region (**Supplementary Fig. 3.2E**).

Collectively, the structural observations, location of cotransin resistance mutations, and molecular dynamics simulations support the proposed binding mode of KZR-8445 within the lumenal cavity of Sec61α, as well as a critical role for N300 in KZR-8445 binding and inhibition.

### Structural insights lead to improved signal peptide selectivity

Previous evidence suggested that cotransin-family Sec61 inhibitors compete with nascent signal peptides for opening the lateral gate and inserting into the lipid bilayer. In particular, signal peptide mutants with increased hydrophobicity or helical propensity were better able to compete, as revealed by a right-shift in the cotransin dose-response curve^16,34^. To gain structural insight into the competitive relationship between nascent signal peptides and cotransins, we superimposed our structure of KZR-8445/Sec61 with two other Sec61 structures, each of which captures a signal peptide inserted between the lateral gate helices in a distinct manner. Comparison of our structure with the prolactin signal peptide bound to mammalian Sec61^6^ reveals a major outward shift of the lateral gate helices TM2, TM3, and TM7 in the KZR-8445 structure (**Fig. 4A**). TM2 packs against the hydrophobic signal peptide, which adopts an alpha-helical conformation and is partially exposed to the lipid bilayer. Strikingly, Sec61 TM2 and the prolactin signal peptide overlap substantially with the KZR-8445 binding site. This analysis indicates that KZR-8445 and the prolactin signal peptide directly compete for binding to a functionally critical site within the Sec61 lateral gate. Compared with most signal peptides tested in the SP-GLuc secretion assay, prolactin is relatively resistant, such that higher concentrations of KZR-8445 are required to inhibit its secretion (IC50 ∼1,000 nM, **Supplementary Fig. 1F**). Hence, a likely contributing feature to Sec61 client selectivity is the relative affinity of KZR-8445 and the nascent signal peptide for overlapping binding sites within the Sec61 lateral gate.

A cryo-EM reconstruction of yeast Sec61 (post-translational translocon with Sec62/63/71/72) bound to the yeast mating type pheromone α-factor^35^ depicts a related but distinct mode of signal peptide binding to the lateral gate (**Fig. 4B**). In contrast to the prolactin signal peptide, which tightly intercalates between TM2 and TM7 and occupies the same region as KZR-8445, the α-factor signal peptide forms a shorter helical segment and docks near the cytosolic ends of TM7 and TM8, which only partially overlaps with the KZR-8445 binding site. Nevertheless, the bulky bromobenzyl-Trp R-5 side chain in our cryo-EM model of KZR-8445 projects into the Sec61 cytosolic vestibule and would likely block many signal peptides from accessing this site (**Fig. 4C**). This observation suggested the possibility of further enhancing signal peptide selectivity by reducing the size of the R-5 side chain, exemplified by KZR-9508 (**Fig. 4D**). We hypothesized that by removing the steric block afforded by the bromobenzyl-Trp side chain, signal peptides entering the cytosolic vestibule would have greater access to the lateral gate. Moreover, the smaller ethyl-Trp side chain in KZR-9508 could potentially impart decreased intrinsic binding affinity for Sec61, facilitating competitive displacement by signal peptides that are otherwise blocked by KZR-8445. We directly compared KZR-8445 and KZR-9508 against a panel of five Sec61 client-GLuc reporters and found that KZR-9508 exhibited greater selectivity (**Fig. 4E**). Whereas KZR-9508 was 10-20 fold less potent than KZR-8445 against 4 out of 5 GLuc reporters, it was nearly equipotent (IC50 ∼100 nM) against the HER3 GLuc reporter. These data suggest that most KZR-8445-sensitive signal peptides, regardless of their precise Sec61 binding mode, are better able to competitively displace KZR-9508. By contrast, a small minority of signal peptides (including HER3) fail to compete with KZR-9508 for binding to the lateral gate (see Discussion).

## DISCUSSION

Many secreted and integral membrane proteins play causal roles in human disease. Preventing their biogenesis by blocking Sec61-dependent entry into the secretory pathway could have profound therapeutic implications. Substrate-nonselective Sec61 inhibitors, including mycolactone and apratoxin, have been described^12^. However, because Sec61-mediated protein import is an essential process in healthy cells, these inhibitors are probably too toxic to be developed as therapeutics^36^. By contrast, substrate-selective Sec61 inhibitors, including cotransin-related cyclic heptadepsipeptides and CADA-related cyclotriazadisulfonamides^13,14,37^ inhibit the biogenesis of a subset of secreted and membrane proteins. How substrate-selective Sec61 inhibitors engage the dynamic translocation channel and discriminate among Sec61 clients in a signal peptide-dependent manner has remained a mystery. By determining the cryo-EM structure of Sec61 bound to the cotransin-related inhibitor KZR-8445, we provide new insights into how substrate-selective Sec61 inhibition can be achieved, as compared to relatively nonselective inhibitors such as mycolactone.

When bound to KZR-8445, the Sec61 lateral gate adopts an open conformation, reminiscent of the open lateral gate observed in the yeast posttranslational Sec61 translocon^27,28^. KZR-8445 binds within the central pore of Sec61 at its lumenal end, a site that maps to previously identified cotransin resistance mutations^16,17^. In addition to contacting the lateral gate where signal peptides ultimately insert into the lipid bilayer, KZR-8445 directly contacts the Sec61 plug helix, which is ordered in our structure, and thereby prevents the Sec61 channel from opening toward the ER lumen. Recently, a cryo-EM reconstruction of Sec61 bound to a substrate-nonselective inhibitor, mycolactone A/B, was described^29^. In contrast to KZR-8445, mycolactone binds to the cytosol-exposed region of the Sec61 lateral gate, far from the lumenal plug domain (**Fig. 5A**). The proposed mycolactone binding site overlaps with the site occupied by the α-factor signal peptide in the yeast post-translational translocon^35^. We speculate that most signal peptides initially traverse this site prior to inserting into the lipid bilayer, which is presumably required to open the lumenal plug and promote translocation of the growing nascent chain. Steric blockade by inhibitors such as mycolactone would thus prevent translocation of a broad range of Sec61 client proteins.

**Figure 5.**
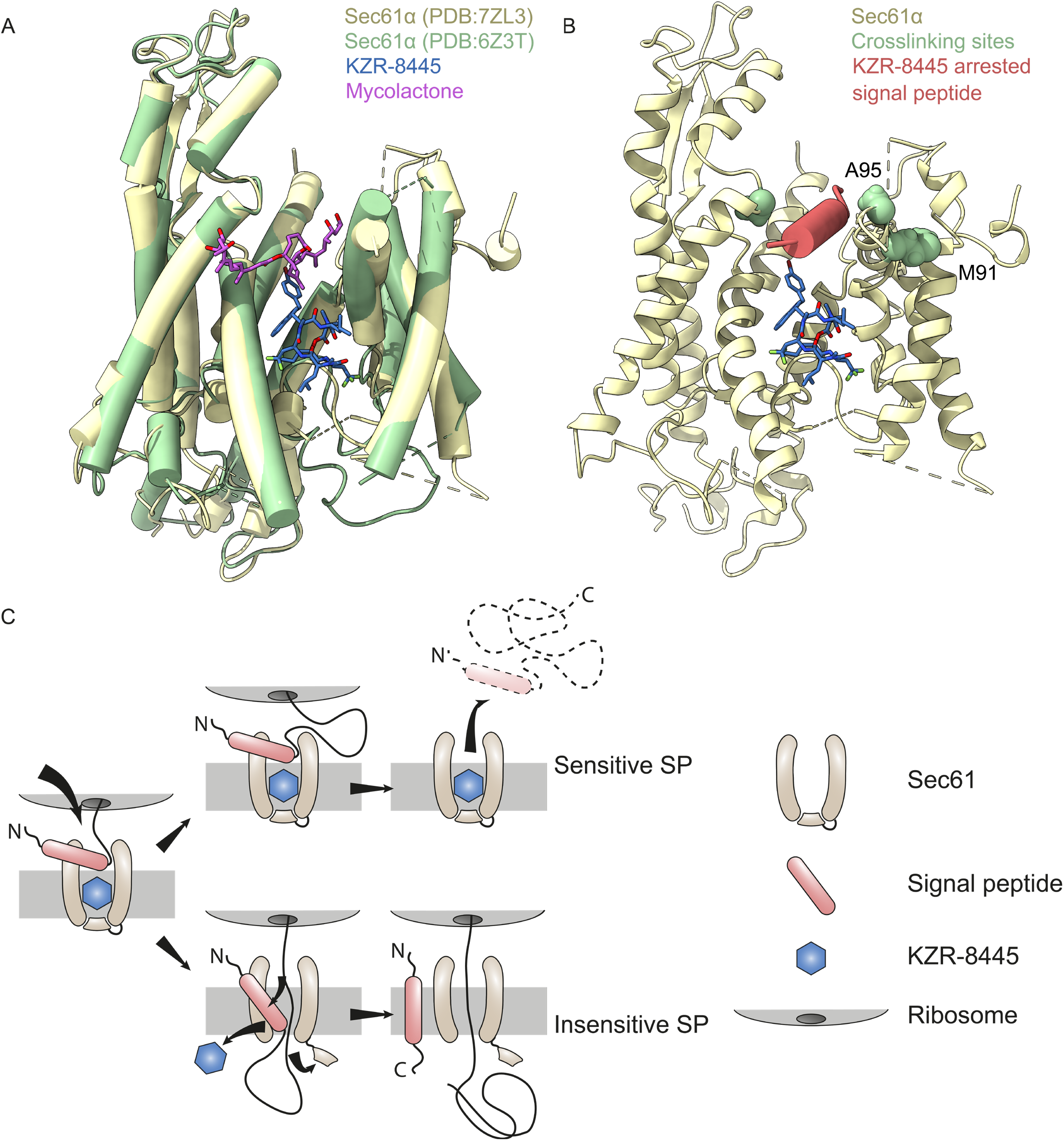
Proposed model for substrate-selective Sec61 inhibition. **(A)** Comparison of KZR-8445 (blue) bound to Sec61α (PDB:7ZL3) and mycolactone (purple) bound to Sec61α (green, PDB:6Z3T). Mycolactone binds Sec61α in a similar conformation as KZR-8445, but at the cytosolic tip of the lateral gate. **(B)** KZR-8445 bound Sec61 (wheat) with residues outlining the cytosolic entrance to the lateral gate which crosslink with TNFα signal peptide in the presence of cotransin highlighted in green. Placement of a putative cotransin arrested signal peptide (red) is guided by proximity of crosslinking residues and occlusion of the channel by KZR-8445. The bulky R-5 group of KZR-8445 projects in the direction of the elongating nascent polypeptide and the overall docking site of signal peptides. **(C)** Substrate-selective inhibitors such as KZR-8445 arrest specific signal peptides in a nonproductive conformation at the cytosolic tip of the lateral gate and in proximity to the KZR-8445 R-5 group, which is important for determining the range of inhibited Sec61 clients. Sensitive signal peptides are unable to progress in the insertion pathway and in cells are displaced in the cytosol. Drug-resistant signal peptides are able to progress further along the insertion pathway. Intercalation between Sec61 lateral gate helices likely leads to inhibitor dissociation and allows complete insertion of the nascent polypeptide into the secretory pathway.

Our structure provides clues into how KZR-8445 achieves substrate-selective Sec61 inhibition and how selectivity can be further enhanced. Superimposition of structures containing a nascent signal peptide bound to Sec61^6,35^ reveals that certain signal peptides (e.g. α-factor) dock in close proximity to the R-5 side chain, whereas others (e.g., prolactin) dock to a site that overlaps extensively with the KZR-8445 binding site. We speculate that in the presence of KZR-8445, a subset of signal peptides stably dock to the mycolactone binding site (**Fig. 5A and 5B**), and fail to promote release of the lumenal plug domain. Such Sec61 clients would ultimately fail to undergo cotranslational translocation into the ER. Our previous work with the cotransin analog CT8 suggests that the N-terminal signal anchor (TMD) of TNFα falls into this category^16^. The open conformation of Sec61 observed in our structure appears to be compatible with an α-helical signal peptide or TMD nestled between KZR-8445 and the lateral gate residues S383, A95 and M91 (**Fig. 5B**). Consistent with this model, these residues were previously shown to reside near the TNFα TMD in the context of a pre-insertion complex stabilized by CT8^16^.

Our finding that KZR-9508 is more selective than KZR-8445 suggests that signal peptide selectivity can be tuned by altering the structure of the R-5 side chain. We propose a model for substrate-selective inhibition in which sensitive signal peptides initially engage Sec61 at the cytosolic tip of the lateral gate (**Fig. 5B**), where they are stabilized in a nonproductive configuration, possibly via direct interactions with a composite surface defined by Sec61 and R-5 of the bound cotransin. This mechanism is reminiscent of nascent chain-selective ribosome inhibitors, which bind to a composite surface defined by the ribosome exit tunnel and specific nascent polypeptide sequences that traverse the exit tunnel^38–40^. Cotransin-resistant signal peptides, on the other hand, can apparently displace the bound inhibitor, intercalate between the lateral gate helices, insert into the lipid bilayer, and ultimately promote opening of the lumenal plug domain.

Secretory proteins such as TNFα, IL-1β and IL-6 act as inflammatory mediators that promote joint damage during progression of arthritis^41^. We demonstrated that KZR-8445 inhibits the stimulated secretion of IL2, TNFα, and GM-CSF in primary mouse splenocytes and human PBMCs. We tested KZR-8445 in a collagen antibody-induced mouse model of rheumatoid arthritis^42^, where it ameliorated clinical arthritis symptoms, likely resulting from selective blockade of several proinflammatory secretory proteins. Unlike previously described Sec61 inhibitors that have been tested in animals, KZR-8445 was well tolerated as evidenced by weight gain with continued dosing, relative to vehicle-treated mice (**Fig 1D**). The observation that KZR-8445 prevents SARS-Cov-2 replication (**Supplementary Fig. 1A–D**), suggests that inhibition of viral glycoprotein biogenesis could potentially be a useful antiviral strategy^19,21^.

Future structural studies of Sec61 bound to defined signal peptides and client-selective cotransins will inform the rational design of small molecules with improved or altered selectivity toward distinct signal peptides and N-terminal transmembrane targeting segments. Potential therapeutic targets include cancer drivers such as HER3, which are currently difficult to target with small molecules^43^, as well as pro-inflammatory cytokines such as TNFα.

## EXPERIMENTAL PROCEDURES

### Mice

BALB/c mice (H-2d) were purchased from Taconic Biosciences. All animal studies were conducted in compliance with the NIH Guide for the Care and Use of Laboratory Animals and approved by the Kezar Life Sciences Institutional Animal Care and Use Committee.

### Arthritis model

Anti-collagen antibody induced arthritis (CAIA) was induced in 7-8 week old female BALB/c mice (kept on breeder chow) by intravenous (IV) administration of 1.75 mg of a cocktail of 5 antibodies against type II collagen (Chondrex, Redmond, WA) followed by intraperitoneal challenge with 25 μg LPS on day 3. Treatment was initiated after clinical signs of arthritis were observed (day 4). Paws were scored for disease severity on a 0 (no disease) – 4 (maximal swelling) scoring system and summed for individual animal scores. Statistical analyses (two-way ANOVA followed by Bonferroni post hoc analysis) was performed using GraphPad Prism Software (version 7.01). Statistical significance was achieved when p was less than 0.05. For efficacy studies, KZR-8445 was formulated in an aqueous solution of 10% ethanol/10% (w/v) Kolliphor EL and was administered three times a week every other day (QODx3) as an IV bolus. Dexamethasone was purchased from Sigma and administered three times a week every other day (QODx3) intraperitoneally.

### Purification and biochemical characterization of the ribosome–Sec61 complex

Sheep pancreatic ER microsomes (SRM) were isolated according to the method described earlier^44,45^. 10 μM KZR-8445 was added to the microsomes and incubated on ice for 30 min. LMNG at a final concentration of 1% was then added to solubilize the microsome for 30 min on ice. Solubilized material was centrifuged at 21000 x g and further purified using 1 ml Superose-12 gel filtration resin in 50 mM HEPES (pH 7.4), 200 mM KoAc, 10 mM MgoAc and 1 mM DTT. The peak fraction containing approximately 400 nM RNC-Sec61 was immediately used for freezing cryo-EM grids. Photo-affinity labeling and click chemistry^18^ were used to confirm binding of KZR-8445 to Sec61.

### Grid preparation and data acquisition

Freshly prepared ribosome–Sec61–KZR-8445 complex at 300–500 nM concentration was applied on glow discharged holey-carbon grids (Quantifoil, R1.2/1.3 with a 2-nm carbon film) and flash frozen with an automated plunge freezer (Leica microsystems). Data sets were collected on a 300 kV FEI Titan Krios TEM (Thermo Fisher Scientific) equipped with a direct electron detector (K3 Summit, Gatan) with GIF Quantum energy filter (Gatan). The images were collected at a dose rate of 0.97 e−/s/Å2 and with an exposure time of 3 s. Movie stacks (50 frames each) were recorded using a super-resolution mode. The magnification was set at 105,000×, resulting in a pixel size of 0.415 Å, and the underfocus ranged from −0.7 to −2.2 μm.

### Image processing

All cryo-EM data processing was performed with Relion 3.046 maintained within the Scipion 3.0.7 software package. A total of 1,089,031 particles were picked from 30,230 motion-corrected micrographs with SPHIRE-crYOLO47, contrast transfer function parameters were estimated using CTFFIND448, and all 2D and 3D classifications and refinements were performed using RELION49. A total of 136,742 selected particles contributing to the best 3D classes were then subjected to iterative rounds of 3D refinement until the FSC converged at 3.5 Å. The output particles from refinement were then 3D classified without alignment, which generated 10 classes with clearly distinguishable translating and non-translating ribosomes. So as to preclude density contributions from nascent polypeptides, only non-translating ribosome-Sec61 complexes were submitted to iterative CTF refinement, resulting in a final map which resolved to 3.2 Å resolution.

### Model building and refinement

An initial model for Sec61 was built using the structure of signal peptide-engaged Sec61 (PDB: 3JC2). The model was used to cut out the ribosome density from a map of non-translating ribosome–Sec61-KZR-8445 particles. The ribosome density was initially removed from the map by using the Map Eraser function in USCF ChimeraX^50^ followed by map truncation outside the Sec61 region using phenix.map_box^51^. The resulting truncated map was then used in automatic molecular dynamics flexible fitting with the program Namdinator^52^. The final model was built using Coot and Phenix real-space refinement^53^. The most likely low energy states of KZR-8445 were calculated as described^30,31^. All figures were generated using USCF ChimeraX^54^.

### Luciferase reporter assay

HEK293T cells were transfected with plasmid encoding a luciferase with the signal peptide of either VCAM or preprolactin (pPL). After trypsinization, cells were diluted to seed a 96-well flat bottom plate with 1.6 × 10^4^ cells per well and allowed to adhere for 6 h. Media was removed, cells were washed once with PBS, and fresh media containing dilutions of either KZR-8445 or mycolactone was added (n=4). After 24 h luciferase-containing media was transferred to a clean 96-well plate and Luciferase activity of the media was estimated using a Gaussia-GLOW Juice Luciferase Assay kit from P.K.J Biotech per the manufacturer’s instructions. Luminescence was measured using an EnSpire Multimode plate reader (PerkinElmer). Data was analysed using GraphPad Prism 8, and calculated IC50 values plotted using the same software.

### Molecular dynamics simulation

The final model of Sec61/KZR-8445 obtained from Coot was used as a starting point for molecular dynamics simulations (MD) with two independent simulation models and simulation engines. For simulations with the NAMD package^55^, the protein and ligand were prepared using the ambertools20 package^56^. The protein and docked ligand were manually embedded in a 100 Å × 100 Å lipid bilayer composed of POPC. The system was solvated with TIP3P water and the system charge balanced with sodium and chloride ions. The Amber14SB57, GAFF2, and LIPID14 forcefields^58^ were employed. The system was gradually equilibrated using a range of harmonic restraint selections and weights. After equilibration was complete, production dynamics of the unrestrained system was conducted with a 2 fs timestep for 1 ns. Analysis of the trajectory yielded a number of candidate conformations of the ligand and local site environment that were compared with the available map. Another set of simulations was performed using the mutually compatible force fields of the CHARMM family. Here, Sec61 was embedded in a multi-component membrane mimicking the ER composition, and five independently generated replicas were simulated for 1 μs. In addition, a 1 μs-long control simulation was performed after removing KZR-8445 from the resolved structure. The binding modes of KZR-8445 were clustered and the clusters analyzed for specific interactions and structural properties. Further details are available in the Extended Data.

## DATA AVAILABILITY

Coordinates of the KZR-8445-bound structure of the mammalian translocon and the corresponding electron density map have been deposited to Protein Data Bank accession code PDB 7ZL3 and Electron Microscopy Data Bank accession code EMD-14776, respectively.

## AUTHOR CONTRIBUTIONS

SR established a protocol for sample preparation for photocrosslinking, cryo-EM, and image processing. VA and JTH planned the image processing and classification strategy. SR, DT, VA, AJ, CK, M Javanainen, M Jacobson, JTH, JT, VOP together solved the structure. EL, DT, JLA, TM carried out the luciferase reporter and other cellular assays. TM performed mouse model studies. SK, NW, TS carried out the SARS-Cov-2 experiments. Chemical synthesis and compound characterization was carried out by PPS, GBC. SR, JT and VOP drafted the manuscript. All authors discussed the results, contributed figures and text, and commented on the manuscript.

## ACKNOWLEDGEMENTS

Research was supported by funding: VOP from the Academy of Finland (338836 and 314672), the Sigrid Juselius Foundation and the Jane and Aatos Erkko Foundation. JTH from the Academy of Finland (314669). M Javanainen from the Academy of Finland (338160) and Emil Aaltonen foundation. JT from UCSF PBBR. We are grateful to Juho Kellosalo for discussion and suggestions on the manuscript. We are thankful to Pasi Laurinmaki and Benita Löflund (CryoEM unit, Institute of Biotechnology, University of Helsinki) for the technical support. We thank Jason van Rooyen (Beamline scientist, Diamond Light Source, UK) for the data collection. We acknowledge CSC – IT Center for Science (Espoo, Finland) for computational resources and Prof. Susan Shao for advice in creating the molecular movies.

