## Supplementation Information for "Signal peptide mimicry primes Sec61 for client-selective inhibition"

**SUPPLEMENTARY RESULTS AND DISCUSSION** 2
**SUPPLEMENTARY FIGURES** 3
**SUPPLEMENTARY TABLE** 15
**SUPPLEMENTARY MATERIALS AND METHODS** 17
**BIOLOGY** 17
Chemicals and reagents 17
Biochemical characterization of ribosome-Sec61 complexes 17
Ribosome-Sec61 complex purification 17
Grid preparation and data acquisition 18
Image processing 18
Protein modeling 19
KZR-8445 modeling 19
Luciferase assay 19
SARS-CoV-2 infections 20
Generation of stable Sec61 mutant cell lines 20
Atomistic molecular dynamics simulations 21
CTG cell viability assays 22
Cytokine secretion assays 23
**CHEMISTRY** 25
Experimental Procedures and Characterization data for KZR-8445 and KZR-9508 25
**REFERENCES** 33

### SUPPLEMENTARY RESULTS AND DISCUSSION

#### KZR-8445 prevents SARS-CoV-2 replication and S-protein biogenesis in cell culture

Inhibiting protein biogenesis at the endoplasmic reticulum has been shown to be effective in preventing replication of different viruses such as influenza, HIV, dengue and recently SARS-CoV-2[1,2]. Therefore, we set out to test whether the moderately substrate-selective KZR-8445 would exhibit SARS-CoV-2 inhibitory properties in cell culture. SARS-CoV-2, the causative agent of the COVID-19 pandemic, is an enveloped, single-stranded, positive-sense RNA virus in the family *Coronaviridae*. SARS-CoV-2 genome is approximately 30 kb long and encodes for 14 open reading frames (ORFs)[2,3]. ORFs 1a and 1b cover the majority of the genome and encode for 16 non-structural proteins required for the viral replication and transcription complexes. The major structural proteins spike (S), membrane (M), envelope (E), and the nucleocapsid (N) are translated from the 3´end of the genome, and are synthesized within the endoplasmic reticulum for virus particle assembly[2]. To assess whether KZR-8445 would inhibit viral particle production, we first measured green monkey kidney (VeroE6) cell viability after wild type SARS-CoV-2 infection and pretreated two hours prior to infection with an increasing dose of KZR-8445. At 48 hours post infection (hpi), cell viability was measured using a luminescence based ATP-assay. KZR-8445 rescued cells from virus-induced cytopathic effect at 1-10 µM concentrations without any observed cytotoxicity **(Supplementary Fig. 1A)**. We noticed that KZR-8445 did not fully prevent release of viral particles into the cell culture medium as assessed by RT-qPCR of the viral RNA (**Supplementary Fig. 1C**). However, evaluation of viral titers in the culture supernatant by TCID50 measurement showed a dramatic dose-dependent decrease in the amount of infectious virus particles (**Supplementary Fig. 1B**), which closely coincided with the effects observed for prevention of virus-induced cytopathy. Reduction in secreted viral particles was also evident by western blotting of cell culture supernatants by antibodies recognizing viral S and N proteins (**Supplementary Fig. 1D**).

### SUPPLEMENTARY FIGURES

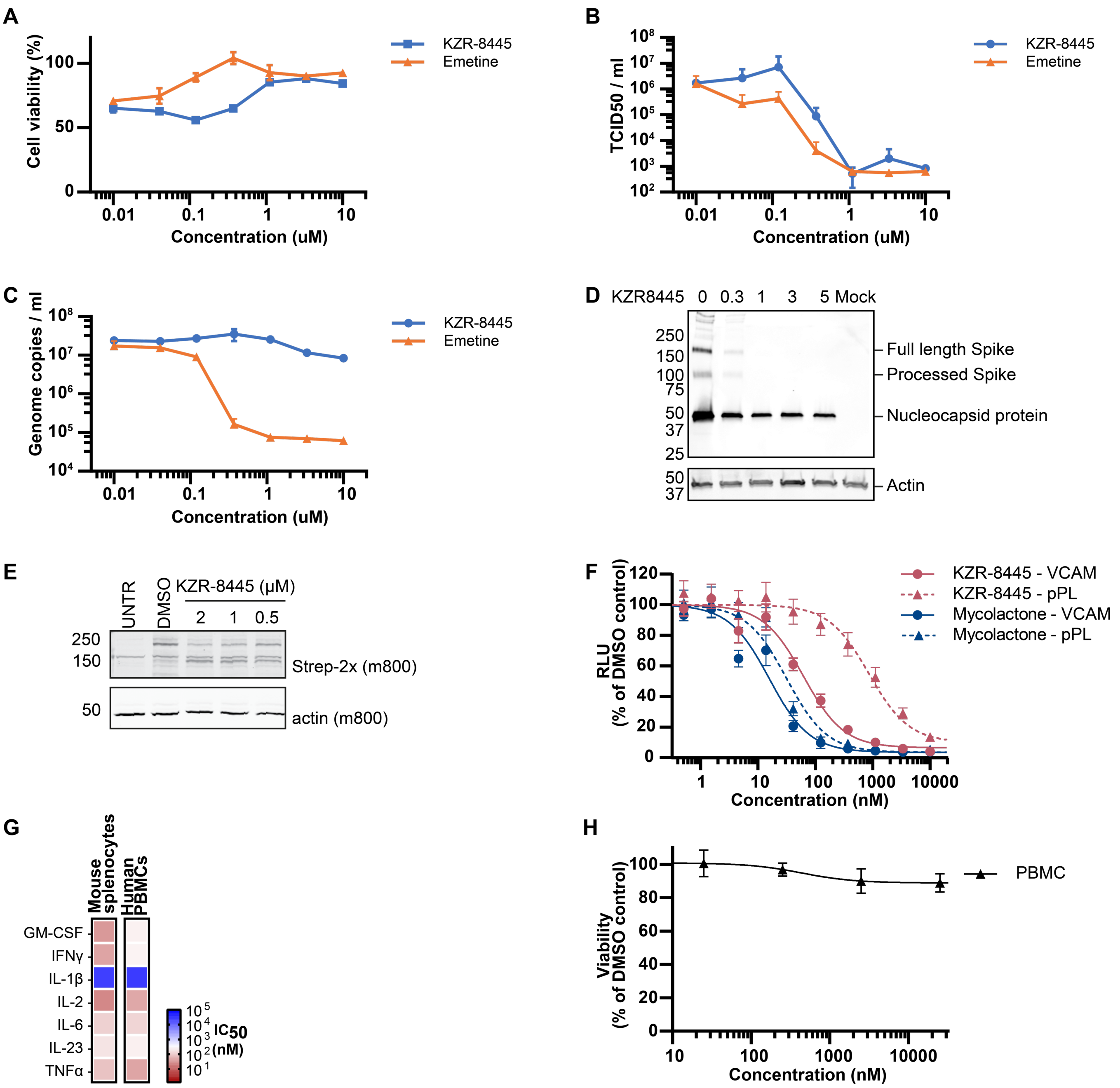


**Supplementary Fig. 1: Cotransin KZR-8445 inhibits SARS-CoV-2 viral replication and biogenesis of the Spike protein. (A)** KZR-8445 rescues virus induced cytopathic effect and **(B)** inhibits infectious virus particle production in VeroE6 cells. Pretreatment of VeroE6 cells with increasing concentrations of KZR-8445 or emetine (translation inhibitor used as a positive control) inhibited the cytopathic effect caused by productive virus replication without loss of cell viability at 48 hours post infection (hpi). In the same concentration range, the infectious virus particle production was reduced by more than 3 logs, when measured by the median tissue culture infectious dose (TCID50) assay from the cell culture supernatant at 48 hpi. **(C)** KZR-8445 does not reduce viral genome copy number in the supernatant of SARS-CoV-2 infected VeroE6 cells. RNA was extracted from the cell culture supernatant of KZR-8445 pretreated cells at 48 hpi and subjected to SARS-CoV-2 RdRp specific quantitative RT-PCR. Despite the inhibition of infectivity and cell death, the viral RNA amount was not reduced. **(D)** KZR-8445 downregulates Spike protein production in infected VeroE6 cells. Cells were pretreated with five concentrations of KZR-8445 (0.3-5μM) and infected with SARS-CoV-2 for 48h. Cell culture supernatants were ultracentrifuged and analyzed for SARS-CoV-2 spike protein expression by Western blotting with anti-RBD (detecting S) and anti-N antibodies (top). Western blot for actin expression in infected cells served as the loading control (below). **(E)** KZR-8445 inhibits production of transiently expressed individual SARS-Cov-2 Spike protein in a dose-dependent manner. Full-length 2xStrep-tagged Spike protein was transiently transfected into HEK293 cells and treated with increasing concentrations of KZR-8445. Spike expression was monitored by Western blotting. UNTR, untransfected control; DMSO, vehicle control. **(F)** KZR-8445 inhibits eGluc2 luciferase secretion in a signal peptide-dependent manner, whereas mycolactone A/B is relatively substrate-nonselective. HEK293T cells were transiently transfected with eGluc2 constructs bearing signal peptides from human VCAM or bovine preprolactin (pPL) and treated with increasing concentrations of KZR-8445 or mycolactone A/B. After 24 h, the amount of secreted luciferase in the medium was determined. **(G)** Effect of KZR-8445 on expression of endogenous pro-inflammatory cytokines in activated human peripheral blood mononuclear cells (PBMCs) or mouse splenocytes. Primary human PBMCs and mouse splenocytes were treated with increasing concentrations of KZR-8445 and either left unstimulated or stimulated with lipopolysaccharide or antibodies against CD3 and CD28. Cells were incubated for 24 h prior to harvesting the supernatant for cytokine analysis via MSD U-PLEX electrochemiluminescent immunoassay. The heat map depicts $IC_{50}$ values derived from dose-response curves for the indicated cytokines. **(H)** KZR-8445 does not significantly affect viability of human PBMCs. Freshly isolated PBMCs were exposed to the indicated concentrations of KZR-8445 for 24 h, and cell viability was evaluated by CellTiter-Glo.

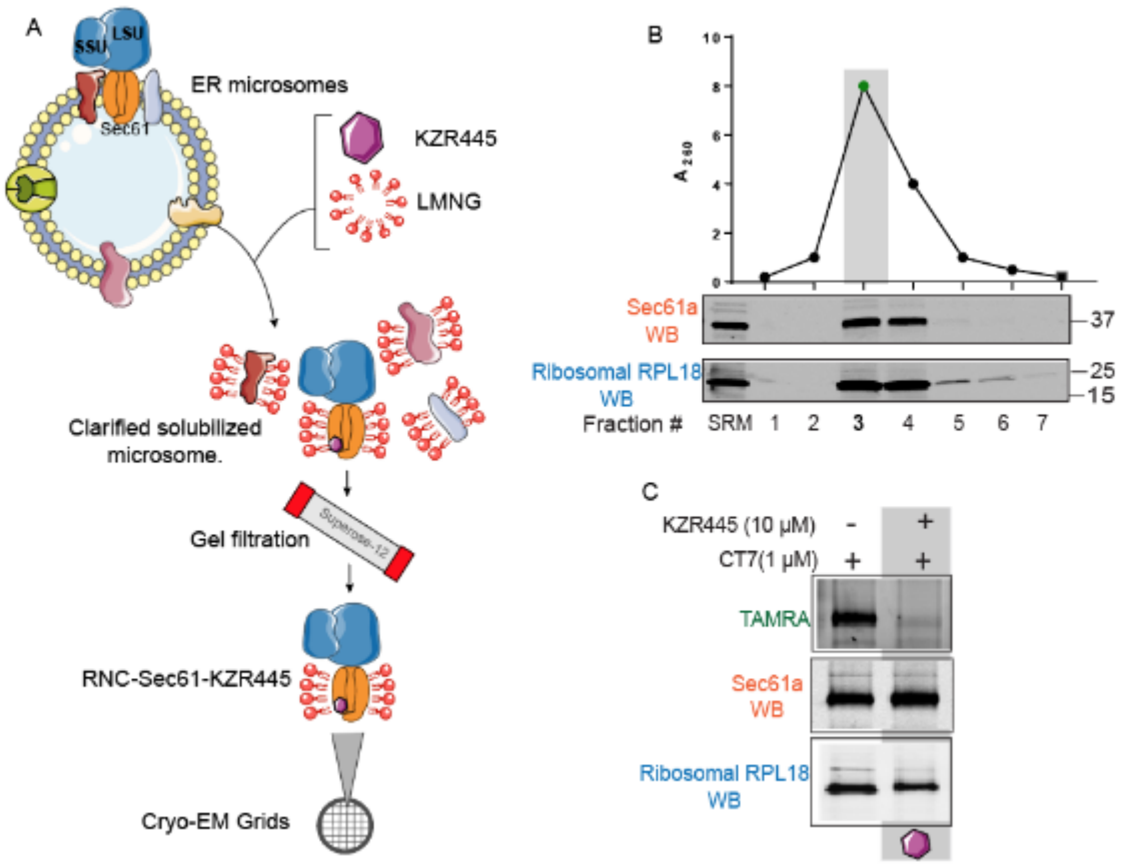


**Supplementary Fig. 2.1: Isolation and biochemical characterization of inhibitor-bound ribosome-Sec61 (RNC) complexes. (A)** Flowchart of purification of KZR-8445-bound RNC-Sec61 complex. Isolated sheep pancreatic rough ER microsomes were supplemented with 1 μM inhibitor and incubated on ice for 30 min followed by solubilization with 1% LMNG for 1 h. Solubilized material was clarified by centrifugation followed by gel filtration **(B)** Gel filtration profile of clarified detergent-solubilized sample fractions with measured absorbance at $A_{260}$. Western blot analysis demonstrates presence of Sec61 and ribosomes in the peak fractions, confirming integrity of the purified complex. **(C)** Photocrosslinking experiment showing competitive binding of KZR-8445 with photo-cotransin (CT7) in the final purified sample. Highlighted sample (gray) was used for cryo-EM analysis.

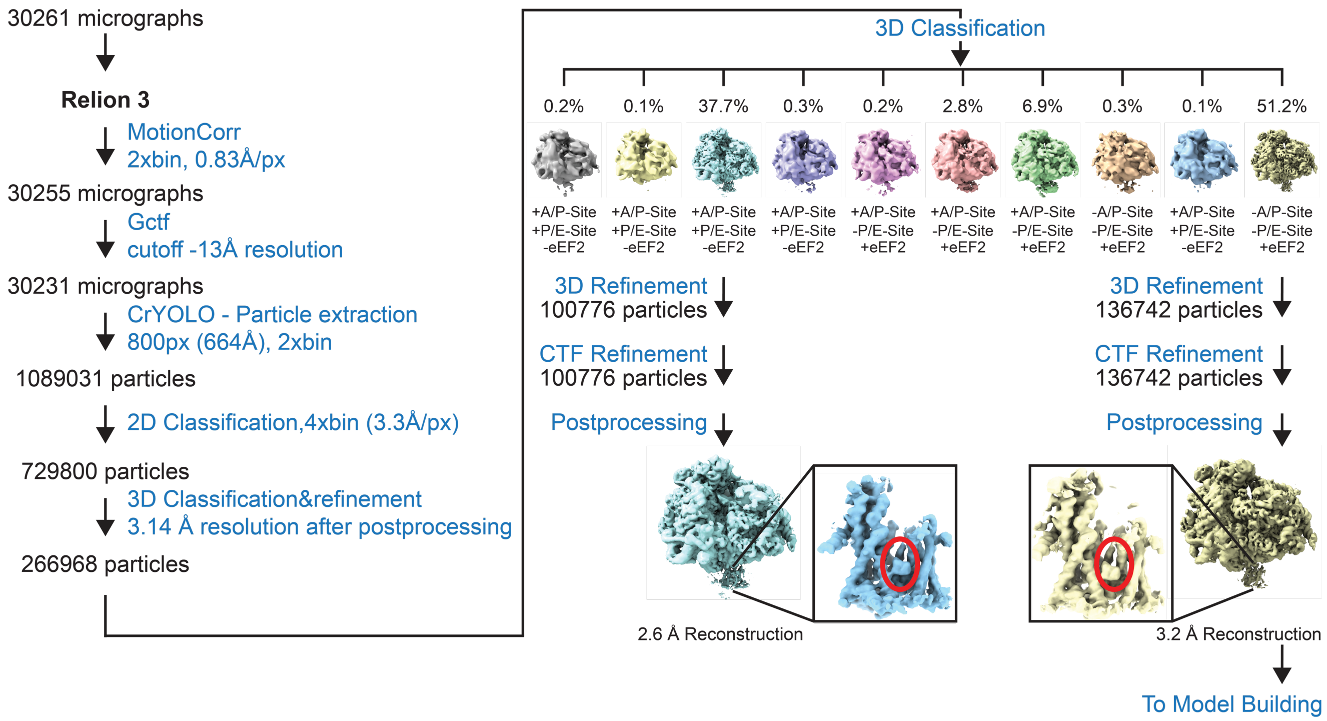


**Supplementary Fig. 2.2: Cryo-EM data-processing workflow for ribosome-Sec61 translocon complexes.**

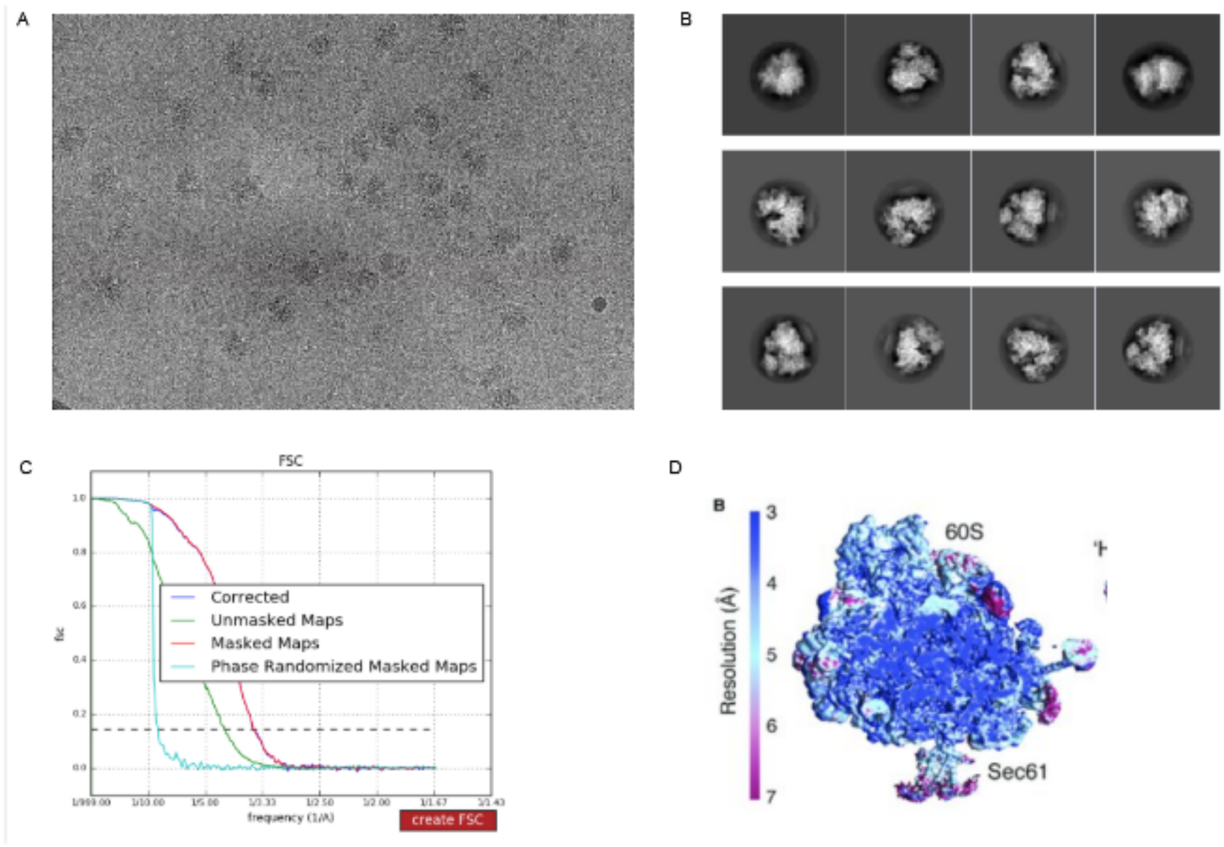


**Supplementary Fig. 2.3: cryo-EM characterization of the ribosome/Sec61/KZR-8445 complex. (A)** A representative micrograph of ribosome-Sec61 complex on holey carbon grids. **(B)** Reference free 2D class averages showing different views of the ribosomes/Sec61 complex. **(C)** Fourier shell correlation (FSC) curves of the final reconstruction map. **(D)** Local resolution of the ribosome/Sec61 complex.

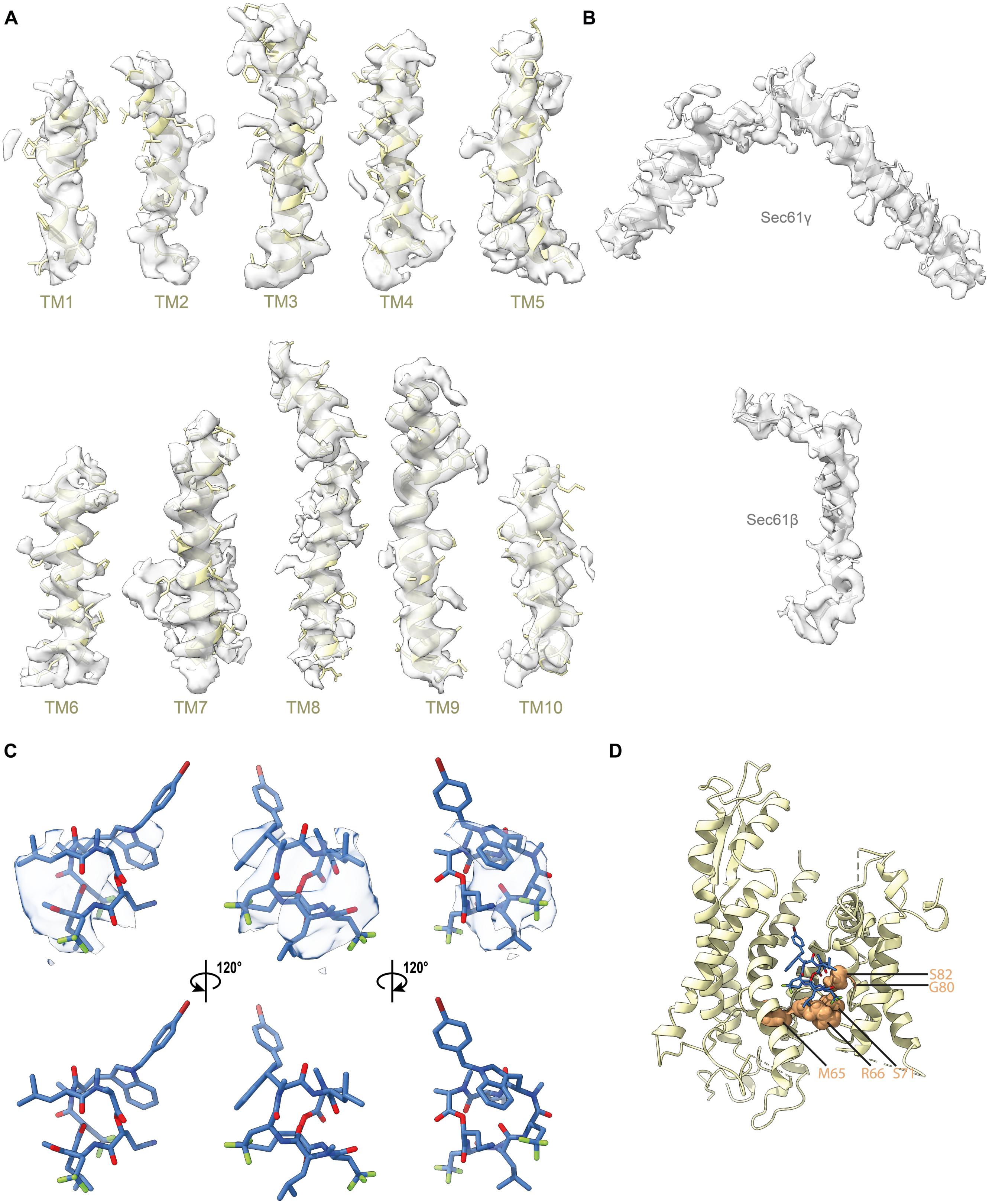
A
B
TM1
TM2
TM3
TM4
TM5
Sec61γ
TM6
TM7
TM8
TM9
TM10
Sec61β
C
D
120°
120°
S82
G80
M65
R66
S71

**Supplementary Fig. 2.4: Structure of the mammalian Sec61 translocon with KZR-8445**

**(A)** Density of the fitted map within 3 Å of each transmembrane helix of Sec61α shown at σ 1.5. **(B)** Density of the fitted map within 3 Å of Sec61γ (top) or Sec61β (bottom), shown at σ 1.5. **(C)** Different views of the density map corresponding to KZR-8445. **(D)** Location of mutations conferring resistance to various cotransins.

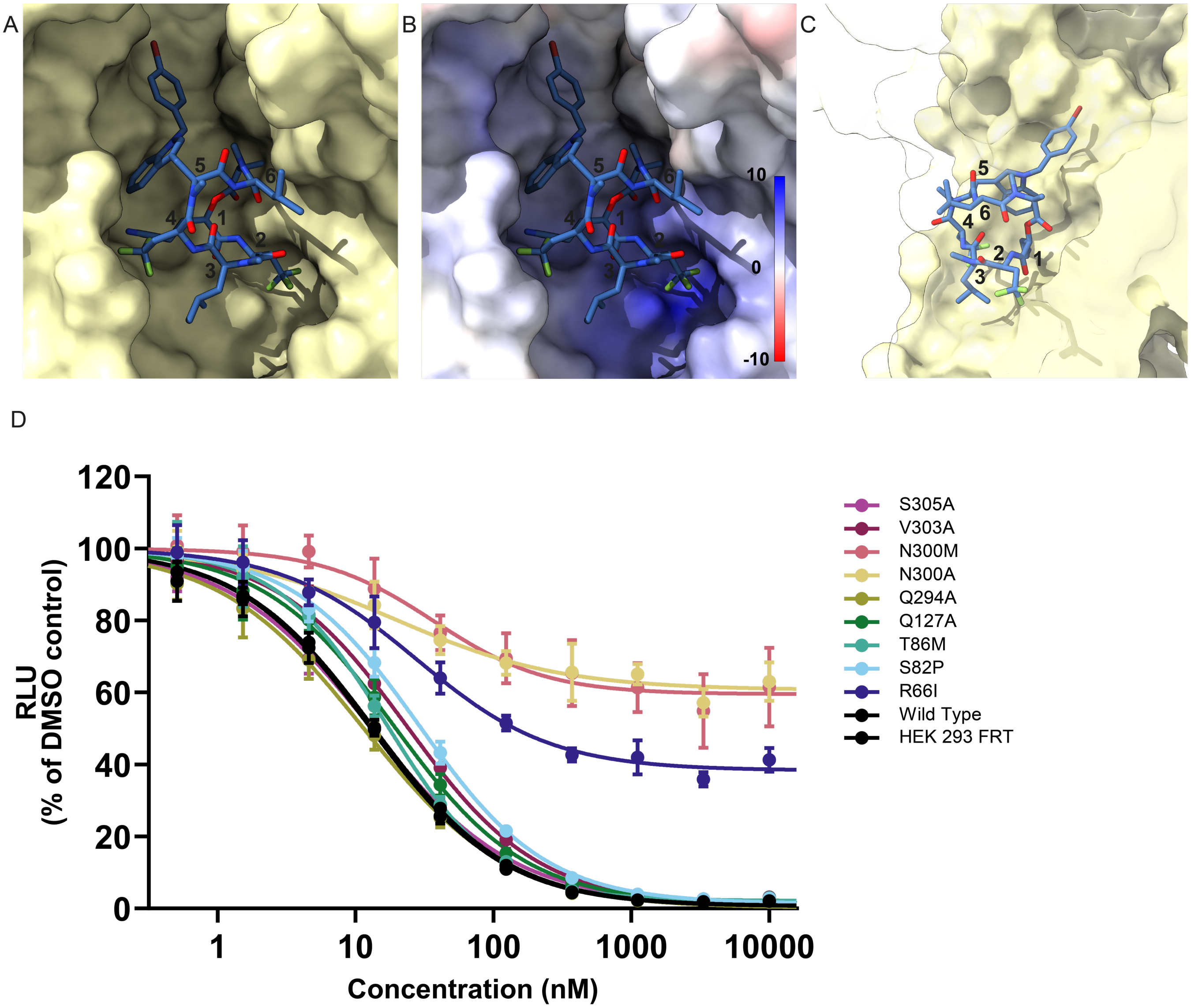


**Supplementary Fig. 3.1. Details of the KZR-8445 binding site.** **(A)** Solvent excluded surface view of the KZR-8445 binding site within Sec61 viewed from the direction of the open lateral gate. **(B)** Same view, but coloured based on the Coulombic potential surface. **(C)** Side view of the solvent excluded surface of the KZR-8445 binding site with intervening surfaces of Sec61α rendered transparent. Part of KZR-8445 is exposed to the lipid bilayer through the open lateral gate. **(D)** KZR-8445 sensitivity of a VCAM-SP GLuc reporter construct in cells expressing the indicated Sec61α mutant.

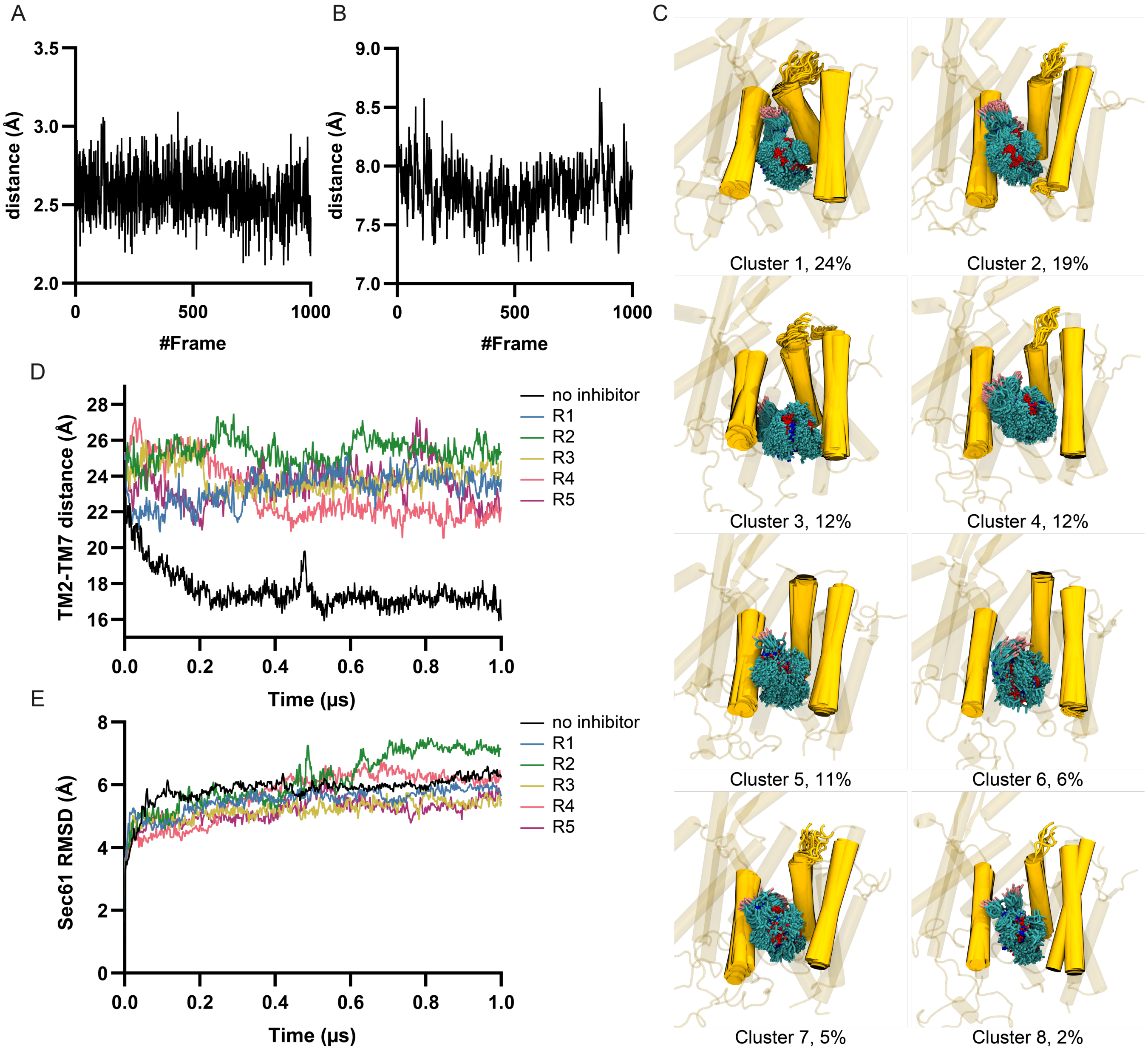


**Supplementary Fig. 3.2: Results from atomistic molecular dynamics simulations.** Simulations were run for either 1 ns **(A** and **B)**, or 1 µs **(C** to **E)**. Data are shown either for different simulated replicas **(D** and **E)** or for clusters extracted from these replicas **(C)**. **(A)** Distance between KZR-8445 nitrile nitrogen to Asn300 side-chain nitrogen over the trajectory. **(B)** Distance between KZR-8445 center of mass and Asn300 center of mass. **(C)** Clustered structures of KZR-8445 bound to Sec61. The 8 structures correspond to 90% of the conformations sampled during the simulations, and the percentage of each cluster represented in the simulation data is indicated. Clustering is based on the conformations of KZR-8445 and the nearby helices TM2, TM3, and TM7 (highlighted; see methods). **(D)** Distance between the centers of mass of the lateral gate helices TM2 and TM7 as a function of simulation time for the five simulation replicas, each 1 µs long. The distances show fluctuations close to the initial value of 2.46 nm observed in the cryo-EM structure, yet the gate remains open in all replicas. In addition, data for a 1 µs-long simulation of Sec61 in the absence of KZR-8445 is shown. In this case, the lateral gate closes rapidly within ~200 ns, after which it remains stable in the closed conformation. **(E)** Root mean squared deviation (RMSD) of the Sec61 structure. All simulated replicas show relatively high RMSD values, indicating the large structural fluctuations of Sec61 due to its two-halved structure. Nevertheless, these values stabilize at around 0.5–0.7 nm, indicating that the protein structures are stable in the simulated membrane.

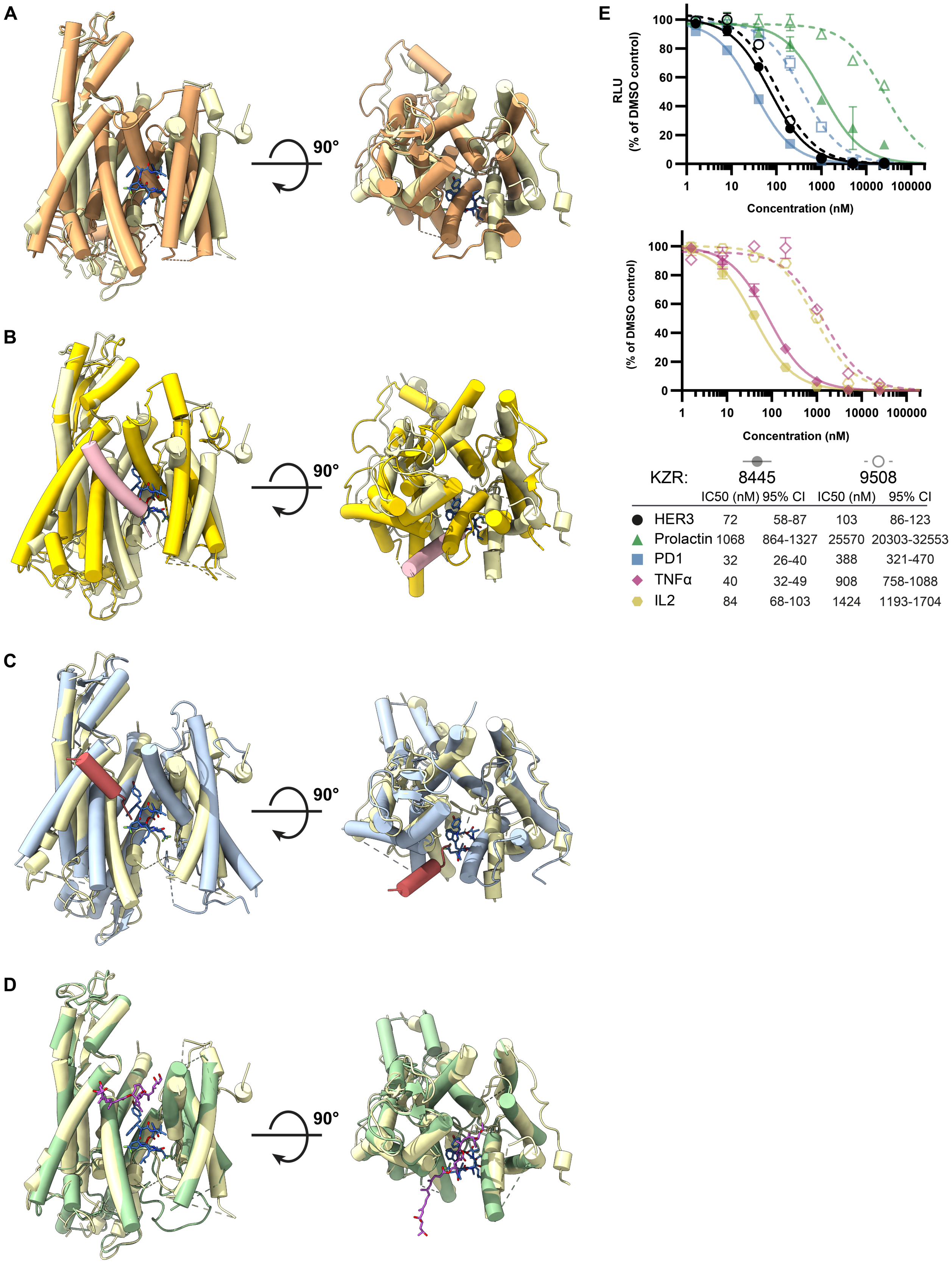


| | IC50 (nM) | 95% CI | IC50 (nM) | 95% CI |
|---|---|---|---|---|
| HER3 | 72 | 58-87 | 103 | 86-123 |
| Prolactin | 1068 | 864-1327 | 25570 | 20303-32553 |
| PD1 | 32 | 26-40 | 388 | 321-470 |
| TNFα | 40 | 32-49 | 908 | 758-1088 |
| IL2 | 84 | 68-103 | 1424 | 1193-1704 |

**Supplementary Fig. 4: Comparison of different structural states of mammalian Sec61. (A-D)** Comparisons of Sec61α (wheat) with bound KZR-8445 (royal blue) and **(A)** “idle” mammalian Sec61α (PDB:3J7Q, orange), **(B)** mammalian Sec61 (PDB:3JC2, yellow) engaged with the preprolactin signal peptide (pink), **(C)** yeast Sec61 (PDB:7AFT, pale blue) engaged with the prepro-α factor signal peptide (red), or **(D)** mammalian Sec61α (PDB:6Z3T, green) with bound mycolactone (purple). **(E)** Cells were stably transfected with dox-inducible Gaussia luciferase (GLuc) reporter constructs fused to the C-terminus of the indicated signal peptides (top), or full-length IL2 or TNFα (bottom). Following treatment with doxycycline and the indicated concentrations of KZR-8445 for 24 h, GLuc activity was quantified.

### SUPPLEMENTARY TABLE

| **Supplementary Table 1. Refinement and Model Statistics** | |
|---|---|
| | **RNC-Sec61-KZR-8445** |
| **Data Collection and Processing** | |
| Magnification | 105000 |
| Voltage | 300 |
| Electron exposure (e-/Å2) | 47,66 |
| Defocus range (µm) | -0.6 to - 2.2 |
| Pixel size (Å) | 0.415 (super resolution) |
| Symmetry imposed | C1 |
| Initial particles images | 1089031 |
| Final particles images | 143690 |
| Map resolution (Å) | 2.6 |
| Fourier Shell Correlation threshold | 0.143 |
| **Refinement** | |
| Map pixel size (Å) | 0.83 |
| Model Resolution (Å) | 2.6 |
| **Model Composition** | |
| Non-hydrogen atoms | 3712 |
| Protein residues | 474 |
| Ligands | KZC:1 |
| Non- | |
| **Root-Mean-Square Deviations (RMSDs)** | |
| Bond lengths (Å) | 0.020 |
| Bond angles (Å) | 2.475 |

| | |
|---|---|
| **Validation** | |
| MolProbity score | 3.06 |
| Clashscore | 23.10 |
| **Ramachandran Plot** | |
| Favoured (%) | 89.69 |
| Allowed (%) | 9.43 |
| Disallowed (%) | 0.88 |

### SUPPLEMENTARY MATERIALS AND METHODS

### BIOLOGY

#### Chemicals and reagents

Lauryl maltose neopentyl glycol detergent was from Anatrace Inc. Superose-12 gel filtration media was obtained from GE healthcare. Anti-Sec61α and anti-RPL16 antibodies were from Abcam. CryoEM grids were purchased from Quantifoil Micro Tools GmbH.

#### Biochemical characterization of ribosome-Sec61 complexes

Sheep rough ER microsomes (SRM) were isolated according to the method described earlier[4,5]. Microsomes were then resuspended in buffer containing 50 mM HEPES (pH 7.4), 200 mM KOAc, 10 mM $Mg(OAc)_2$ and 1 mM DTT and treated with micrococcal nuclease in the presence of 1 mM $CaCl_2$ in order to convert the polysomes into monosomes. The reaction was stopped by chelating the $Ca^{2+}$ with 2 mM EGTA. Microsomes were aliquoted and stored at -70°C until further use.

To identify optimal conditions for reconstituting and purifying inhibitor-bound Sec61, photo-affinity labeling and click chemistry were used as previously described[6,7]. Detergent-solubilized ER microsomes containing 100 nM Sec61 were first incubated for 30 min with KZR-8445 or DMSO, then with 1 µM photo-cotransin CT7 for 10 min, and cross-linking was performed by UV-irradiation for 10 min. After denaturation with 1% SDS, copper-catalyzed click chemistry was used to label the cross-linked adducts with the TAMRA fluorophore. The labeled proteins were analyzed by SDS-PAGE and in-gel fluorescence followed by Western blotting using anti-Sec61α and anti-RPL18 antibodies. Using this method, several detergents and solubilization conditions were screened to find the optimal conditions where the inhibitor stably remains bound.

#### Ribosome-Sec61 complex purification

For the purification of inhibitor-bound ribosome-Sec61 complexes, 50 µl of SRM were thawed and KZR-8445 was added to a final inhibitor concentration of 10 µM and incubated on ice for 30min. 1% LMNG was added and the sample was further incubated on ice for 1 hour with occasional mixing. Insoluble material was separated by centrifugation at 22000 rpm for 15 min. The clarified supernatant was loaded on a 1 ml Superose-12 gel filtration column pre-equilibrated with a buffer containing 0.003% LMNG and 1 µM KZR-8445. 10 fractions each containing approximately 100 µl sample were collected and $A_{260}$ absorbance was measured using a nanodrop spectrophotometer. The final concentration of the sample was estimated using the molar extinction coefficient of eukaryotic ribosomes[8]. The peak fraction was supplemented to 10 µM KZR-8445 and incubated for 30 min on ice. The sample was centrifuged at 22000 rpm for 10 min to exclude any aggregates before freezing grids.

**Grid preparation and data acquisition**

Holey-carbon grids (Quantifoil, R1.2/1.3 with 2 nm C) were coated with pentylamine (Sigma aldrich, cat no. 171409) using a method described earlier[9] and glow-discharged with a plasma cleaner. 3 µl of freshly prepared sample at a final concentration of 300-500nM was applied to the grid and blotted for 1.5 sec before vitrification in liquid ethane pre-cooled by liquid nitrogen. Cryo-grid preparation was assisted by an automated plunge freezer (Leica microsystems) with the inner chamber set at 15 °C and 90% humidity. The cryo-grids were prescreened with a 200 kV FEI Talos Arctica microscope (FEI Falcon II camera). Final high resolution data sets were collected on a 300 kV FEI Titan Krios TEM (Gatan K3 summit camera) with GIF Quantum energy filter (Gatan). The images were collected at a dose rate of 0.97 $e^{-}/s/Å^2$ and with an exposure time of 3 s. Movie stacks (50 frames each) were recorded under super resolution conditions. The magnification was set at ×105,000 and the defocus ranged from −0.7 to −2.2 µm. Statistics for data collection are summarized in Supplementary Table 1.

**Image processing**

All cryo-EM data processing was performed with Relion 3.0[10] maintained within the Scipion software package. Frames from 30,261 micrographs were aligned and motion corrected with MotionCor2 using 15 (5x5) patches, the default B-factor of 150 and 2x binning. Defocus values were calculated from the non-dose-weighted micrographs with Gctf [11] using information up to a resolution limit of 10 Å. A total of 31 micrographs with Gctf estimated maximum resolution worse than 20 Å were discarded. A total of 1,089,031 particles were picked from the resulting 30,230 micrographs with SPHIRE-crYOLO[12] using a confidence threshold of 0.05 and box size of 800 pixels. Selected particles were extracted with a binned pixel size of 3.32 Å and 2D classified to remove aberrant particles. Three rounds of 2D classification were carried out, and 30 good classes containing 729800 particles were selected after the final round. These were then refined to generate initial 3D reconstruction with a resolution of 6.7 Å. Alignment information from well-resolved 3D classes containing 266968 particles was used to re-extract the particles with an unbinned super resolution pixel size of 0.83 Å. These particles were then subjected to iterative rounds of 3D refinement and CTF refinement until the FSC converged at 3.6 Å.The output particles from refinement were then 3D classified without alignment that generated 10 classes. Non-translating ribosomes were distinguished from translating ribosomes by A/P/E site occupancy. The final map from non-translating ribosome-Sec61 complexes (containing 136742 particles) was then post-processed to 3.2Å.

**Protein modeling**

Initial model for Sec61 bound to KZR-8445 was obtained by cutting out the ribosome density from a map of non-translating ribosome/Sec61 particles obtained after membrane focused classification. The ribosome density was initially removed from the map by using the Map Eraser function in ChimeraX[13] followed by map truncation outside the Sec61 region using phenix.map_box[14]. The resulting truncated map was then used in automatic molecular dynamics flexible fitting with the program Namdinator[15]. The final model was built using Coot[16]. All figures were generated using ChimeraX[17].

**KZR-8445 modeling**

Initial low energy conformations of KZR-8445 were calculated using the LowModeMD Search method[18] implemented within MOE[19] and as previously described[20,21].These initial models were fitted into the cryo-EM density and refined manually and using real space refinement in phenix.refine.

**Gaussia Luciferase assay**

0.3 x $10^6$ HEK293T cells were seeded per well in a 6-well plate and incubated at 37 °C, 5% CO2, for 24 hours. Each of two wells was transfected with 1 µg of plasmid DNA (encoding Gaussia luciferase fused to the C-terminus of the signal peptide of VCAM or pPL) using PEI in a 5:1 ratio, after which the cells were returned to the incubator for a further 24 hours. Transfected cells were then resuspended following trypsinization, cell count estimated using a TC20 Automated Cell Counter (BioRad) and diluted to seed a 96-well flat bottom plate with 1.6 x $10^4$ cells per well. Cells were returned to the incubator for 6 hours to adhere, after which media was removed, cells were washed once with PBS, and fresh media containing dilutions of either KZR-8445 or mycolactone A/B were added (n=4). Cells were then incubated for a further 24 hours, after which the luciferase-containing media was transferred to a clean 96-well plate and stored at -20°C. Luciferase activity of the media was estimated using a

Gaussia-GLOW Juice Luciferase Assay kit from P.K.J Biotech per the manufacturer's instructions. Luminescence was measured using an EnSpire Multimode plate reader (PerkinElmer). Background luminescence from media placed in a well with neither cells nor drug was subtracted from all values, and measurements calculated as a percentage of luminescence intensity of a well expressing the appropriate luciferase construct in the absence of any drug. Curves were fitted and IC50 values calculated using GraphPad Prism 8.

**SARS-CoV-2 infections**

VeroE6 cells (ATCC® CRL-1586) were cultured in Minimum Essential Media supplemented with 10% fetal bovine serum (FBS; standard FBS hereafter) (Gibco), 2 mM L-glutamine, 100 IU/ml of penicillin and 100 µg/ml of streptomycin, and plated on either 96-well plate (Perkin Elmer) (30 000 cells per well) or on a 6-well plate (200 000 cells per well) for experiments. For viability assay, cells were treated with seven concentrations of KZR-8445 in triplicates for 2 hours at 37 °C and 5% $CO_2$ prior to infection, after which the cells were infected with SARS-CoV-2[22] or mock infected with a multiplicity of infection (MOI) of 0.03 for 48 hours. Supernatant samples were collected and cell viability assay was performed in a 96-well plate using CellTiter Glo 2.0 cell viability assay (Promega) and Hidex Sense reader (Hidex) in a BSL-3 facility. Infectious virus amount was determined by end-point titration assay in quadruplets of each replicate, and presented as $TCID_{50}$/ml[23]. Shortly, 10-fold dilutions of the samples were inoculated to VeroE6 cells, incubated for 5 days, fixed with 10% formaldehyde for 30 min at RT and stained with crystal violet. For RT-PCR, RNA was extracted from cell culture supernatant using QIAamp Viral RNA Mini Kit (Qiagen) and quantitative RT-PCR was performed using primers and probe specific to SARS-CoV-2 RdRP, as described[24]. Emetine was used as a control as it has been described to inhibit SARS-CoV-2 translation[25]. For protein analysis, cells were treated with five concentrations of KZR-8445 2 hours prior to infection. At 48 hours post infection, cell samples were collected in Laemmli sample buffer (LSB) (Sigma) and supernatants were ultra-centrifuged through a 30% sucrose cushion at 141 000 g for 90 min and resuspended in LSB. Both sample types were subjected to SDS-PAGE in 4–15% Mini-PROTEAN® TGX™ gels (Biorad) and transferred onto nitrocellulose membranes. SARS-CoV-2 spike and nucleocapsid proteins were visualized by rabbit anti-RBD (1 µg/ml) and anti-N (200 ng/ml) primary[26], and IRDye® 800CW Goat anti-Rabbit IgG and IRDye® 680CW Goat anti-Rabbit IgG secondary antibodies (Li-Cor).

**Generation of stable Sec61 mutant cell lines**

Wild type dog Sec61 was cloned into pcDNA5/FRT/TO (Thermo Fisher Scientific) and point mutations were generated using PCR. All mutants were sequence verified. HEK293 Flp-In T-REx cells (R78007, Thermo Fisher Scientific) were cultured in Dulbecco's Modified Eagle's Medium (Gibco) supplemented with 10% FBS (Gibco) at 37 °C in a humidified 5% $CO_2$ atmosphere. Approximately 0.3 µg of pOG44 and 1 µg of targeting construct plasmids were mixed in 250 µl of OptiMEM (51985-026, Thermo Fisher Scientific). 3 µl (3 µg) Lipofectamine3000 transfection reagent was then added to the DNA mix and incubated at room temperature for 20 minutes and the whole mixture was added to the cells. Cells were exposed to 50 µg/ml hygromycin B (Invitrogen) and 10 µg/ml blasticidin for 3-4 weeks until the appearance of resistant colonies. Colonies were expanded into T75 flasks and induced with doxycycline (D9891, Sigma) at a concentration of 1–5 µg/ml for 48 hours. Cells were harvested and total RNA was extracted followed by cDNA amplification and sequencing of Sec61 with gene specific PCR primers. Cells were then used for the luciferase reporter assays as above.

**Atomistic molecular dynamics simulations**

The Sec61 structure with the modeled KZR-8445 inhibitor was embedded in a multi-component lipid membrane consisting of 54% POPC, 21% POPE, 10% POPI, 4% POPS, 4% PSM, and 7% cholesterol, mimicking the known ER membrane lipid composition[27–30]. The membrane was solvated in a 0.14 M KCl solution, and additional counter ions were included to neutralize the system. The simulation system contained 800 lipid molecules (400 per leaflet) and 64000 water molecules (80 per lipid) for a total of ~302,000 atoms, and spanned dimensions of 16.2 nm ×16.2 nm ×11.1 nm. Five independent starting configurations were generated using CHARMM-GUI[31,32]. Additionally, we performed a control simulation without KZR-8445 present. The KZR-8445 parametrization was also performed using CHARMM-GUI[33]. The mutually compatible force fields of the CHARMM family were used; CHARMM36m for the protein[34,35] , CHARMM36 for the lipids[36], CGenFF for the inhibitor[37], and CHARMM-specific TIP3P for water[38,39]. The force fields were downloaded from CHARMM-GUI in GROMACS-compatible formats[40]. All six systems were first subject to minimization and equilibration protocols[40], after which each was simulated for 1 µs. and GROMACS 2021 was used to perform all simulations[41].

We used the simulation parameters recommended for CHARMM force fields in GROMACS[40]; The time integration was performed using a leap-frog algorithm with a time step of 2 fs. Buffered Verlet lists were used to keep track of all neighbours[42]. The Lennard-Jones forces were switched to zero between 1.0 and a cut-off distance of 1.2 nm. Long-range electrostatic interactions were included by the smooth particle mesh Ewald algorithm[43,44]. Temperatures of the protein (including the inhibitor), the lipids, and the solvent (water and ions) were separately coupled to a Nosé–Hoover thermostat[45,46] with a target temperature of 310 K and a relaxation time of 1 ps. The pressure was maintained at 1 bar with a Parrinello–Rahman barostat[47]. To maintain a square membrane patch and to respect the symmetry of the system, a semi-isotropic coupling was used with the pressures normal to (*z* dimension) and along the plane of the bilayer (*x* and *y* dimensions) coupled separately. The target pressure was set to 1 bar, the compressibility to $4.5 \times 10^{-5}$ $bar^{-1}$ and the relaxation time constant 5 ps. The bonds involving hydrogen atoms were constrained using p-LINCS[48,49].

The five 1 µs-long simulation trajectories were subject to analyses in two main ways. First, the behavior of each independent replica as a function of simulation time was considered. From these data, we extracted the distance of the lateral gate helices 2 and 7, which describes the openness of the lateral gate. This was achieved using the *gmx distance* tool bundled with GROMACS[41]. Additionally, *gmx rms* was used to extract the root mean squared deviation of the protein structure, which is used to confirm the stability of the protein.

Secondly, we divided the accumulated trajectory based on the conformations of the KZR-8445 and the backbones of the nearby TM helices 2, 3, and 7. One of the five replica simulations was omitted from this analysis, since a cholesterol molecule penetrated to the Sec61 channel through the lateral gate maintained open by KZR-8445 and thus affected the binding pose and the conformations of the nearby helices. This binding significantly altered the conformations of helices 2, 3, and 7. The conformations from the other four 1 µs-long simulations were clustered using the GROMOS algorithm[50] with a cut-off of 0.22 nm. This resulted in 8 clusters that accounted for 90% of all sampled conformations. These clusters were subject to analyses to describe the conformational flexibility of KZR-8445 and the nearby protein helices. The root mean squared deviations were calculated for the KZR-8445 and Sec61 using *gmx rms*. The snapshots were rendered using the tachyon renderer in VMD[51].

#### CTG cell viability assays

Peripheral blood mononuclear cells (PBMC) were isolated from fresh whole blood (AllCells, Alameda, CA) via Leucosep tube (Greiner Bio-One, Monroe, NC), including red blood cell lysis using Pharm Lyse solution (BD Biosciences, Franklin Lakes, NJ). For viability experiments, unstimulated PBMCs were plated at 200,000 cells/well in 100 µl growth media (RPMI 1640 supplemented with 5% FBS, 2 mM L-glutamine, 10 mM HEPES, and 100 IU penicillin/100 µg/ml streptomycin) in 96-well clear polystyrene round-bottom tissue culture-treated plates. 50 µl of growth media and 50 µl of 4x compound stock solution (7-point log dilutions, final concentration range of 0.025-25000 nM, 0.25% DMSO) were immediately added, and the cells cultured at 37 °C with 5% $CO_2$ for 24 hours. Subsequently, plates were centrifuged at ~500xg for 5 minutes at room temperature, and 100 µl of supernatant was removed for subsequent cytokine analysis. For viability determination, 100 µl of CellTiter-Glo (Promega, Madison, WI) was added to the remaining 100 µl cells or growth media. Plates were shaken for 2 min, and the reaction transferred to a black-wall, clear flat-bottom 96-well polystyrene plate for assay readout. The assay plate was allowed to sit for 10 min at room temperature before measuring luminescence using a M1000 Pro plate reader (Tecan, Morrisville, NC).

**Cytokine secretion assays**

For cytokine experiments, human PBMCs and mouse splenocytes were either left unstimulated or stimulated with lipopolysaccharide or antibodies against CD3 and CD28. PBMCs were isolated as above; splenocytes were obtained from 8-12 week old female BALB/c mice. Spleens were harvested into cold PBS supplemented with 100 IU penicillin, 100 µg/mL streptomycin, then kept on ice until tissue disruption/cell isolation was carried out using a 100 µm nylon cell strainer (BD Biosciences, Franklin Lakes, NJ). Strained tissue was centrifuged at ~300xg for 5 min at 4°C, then red blood cells were lysed in Pharm Lyse solution (BD Bioscience, Franklin Lakes, NJ), before being rinsed in PBS and resuspended in growth media (RPMI 1640 supplemented with 10% FBS, 1mM sodium pyruvate, 100 IU penicillin, 100 µg/ml streptomycin, and 0.05 mM beta-mercaptoethanol).

For anti-CD3/CD28 stimulation, 96-well clear polystyrene round-bottom tissue culture-treated plates were coated overnight at 4 °C with 100 µl/well of 2 µg/ml mouse anti-human CD3e (ThermoFisher #MA1-10176, clone OKT3; for PBMCs) or 5 µg/ml hamster anti-mouse CD3e (BD #553057, clone 145-2C11; for splenocytes). Immediately prior to cell plating, anti-CD3-coated wells were emptied and washed twice with 200 µl PBS. PBMCs and splenocytes were plated at a density of 200,000 cells/well in 100 µl of their respective growth media. 50 µl of growth media (for unstimulated cells), 50 µl of 4x LPS (for LPS-stimulated cells; Sigma #L431; final concentration 1 µg/ml for PBMCs, 5 µg/ml for splenocytes), or 50 µl of 4x anti-CD28 (for anti-CD3/CD28-stimulated cells; final concentration 2 µg/ml for PBMCs, 5µg/ml for splenocytes) were immediately added to cells, along with 50 µl of 4x compound stock (7-point log dilutions, final concentration range 0.025-25000nM, 0.25% DMSO). Mouse anti-human CD28 (for PBMCs) was from BD #555725, clone CD28.2; hamster anti-mouse CD28 (for splenocytes) was from BD #553294, clone 37.51. Cells were cultured as above for 24 hours, and supernatant collected for cytokine analysis.

Cytokine concentrations in 100 µl media supernatant were quantified using a MSD U-PLEX electrochemiluminescent immunoassay (Meso Scale Diagnostics, Rockville, MD; #K15067L for human, #K15069L for mouse). Biomarker assays were custom 96-well 7-plex plates for simultaneous analysis of species-specific GM-CSF, IFNg, IL-1b, IL-2, IL-6, IL-23, and TNFa. PBMC supernatant was diluted 1:10 in assay diluent prior to analysis; splenocyte supernatant was assayed neat or diluted 1:4 or 1:10. Assays (including calibrator standard curve generation) were performed according to the manufacturer's instructions and were read on a MESO QuickPlex SQ 120 imager. Compound $IC_{50}$s were calculated using 4-parameter logistic regression of DMSO-normalized dose response curves; LPS-stimulation conditions were used for calculation of IL-1b, IL-6, and IL-23 $IC_{50}$s; anti-CD3/CD28-stimulation for GM-CSF, IFNg, IL-2, and TNFα $IC_{50}$ values.

### CHEMISTRY

#### Experimental Procedures and Characterization data for KZR-8445 and KZR-9508

NMR spectra were recorded on a Bruker spectrometer at 400 MHz. Chemical shifts were reported as parts per million (ppm) from an internal tetramethylsilane standard or solvent references. LCMS was performed on a Waters Alliance HT LC/MS (0.2 mL/min) S8 using an Xterra MS C18 column (Waters) and a water/acetonitrile gradient (0.1% formic acid). Preparative HPLC was performed on a Waters 2545 binary gradient module with a Waters 2998 photodiode array module and using an Atlantis T3 Prep ODB 5 µM 19 × 250 mm C18 column. Analytical thin-layer chromatography was performed with silica gel 60 F254 glass plates (EM Science). Silica gel chromatography was performed with 230-400 mesh silica gel. Flash chromatography was performed using a Teledyne ISCO combiFlash Rf system. All solvents were of ACS grade (Fisher Scientific) and used without further purification. Commercially available reagents were used without further purification.

#### $N^{\alpha}$-(*tert*-Butoxycarbonyl)-$N^{\alpha}$-methyl-L-tryptophan

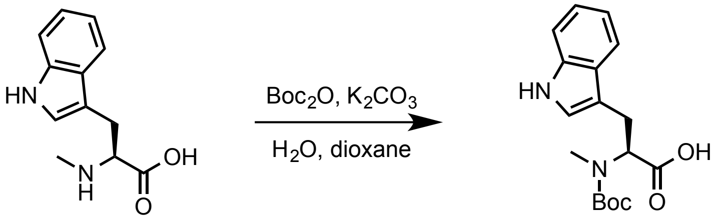


To a suspension of abrine (2.00 g, 9.16 mmol) in water (17 mL) and dioxane (17 mL) was added $K_2CO_3$ (3.60 g, 26.12 mmol) and then, dropwise, a solution of $Boc_2O$ (2.40 g, 11.00 mmol) in dioxane (10 mL). The mixture was stirred at room temperature for 12 h then concentrated to remove dioxane. The reaction mixture was washed with hexane (2 × 20 mL) then acidified to pH ~ 3 with citric acid (10% aqueous solution) and extracted with EtOAc (2 × 20 mL). The combined organic phase was washed with water (10 mL), brine (10 mL), then dried with ($Na_2SO_4$), filtered and concentrated under reduced pressure to afford $N^{\alpha}$-(*tert*-butoxycarbonyl)-$N^{\alpha}$-methyl-L-tryptophan (2.76 g, 95%) as a colorless solid. **$^1$H-NMR** (400 MHz; $CDCl_3$, 2 rotamers): δ 8.14 (broad m, 1H), 7.64 (d, *J* = 7.8 Hz, 1H), 7.39 (d, *J* = 8.0 Hz, 1H), 7.23 (dd, *J* = 7.9, 7.2 Hz, 1H), 7.16 (t, *J* = 7.4 Hz, 1H), 7.09 (s, 0.5), 7.04 (s, 0.5H), 4.91-4.86 (m, 1H), 3.53-3.41 (m, 1H), 3.41-3.36 (m, 0.5H), 3.24-3.17 (m, 0.5H), 2.83 (s, 1.6 H), 2.74 (s, 1.4 H), 1.47 (s, 4.5H), 1.24 (s, 4.5H). **HRMS** (ESI): Calculated for $C_{17}H_{21}N_2O_4$ $[M-H]^-$, 317.1507; Found, 317.1529.

**1-(4-Bromobenzyl)-N$^{\alpha}$-(*tert*-butoxycarbonyl)-N$^{\alpha}$-methyl-L-tryptophan**

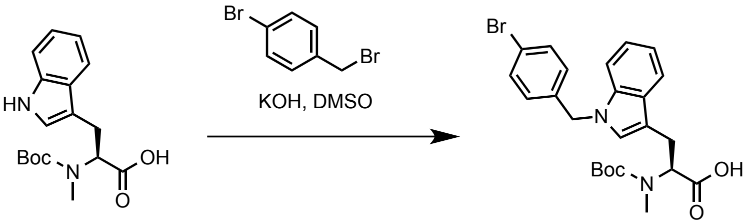


To a stirred solution of freshly powdered potassium hydroxide (4.32 g, 75.38 mmol) in dimethyl sulfoxide (anhydrous, 37 mL) at room temperature under argon was added N$^{\alpha}$-(*tert*-butoxycarbonyl)-N$^{\alpha}$-methyl-L-tryptophan (6.00 g, 18.85 mmol) and the mixture was stirred for 1 h. 4-Bromobenzyl bromide (5.18 g, 20.73 mmol) was then added and the mixture was stirred under argon for 16 h. The solution was diluted with water (10 mL), washed with diethyl ether (2 × 5 mL), and then acidified with citric acid (10% aqueous solution) until pH ~ 3. The mixture was extracted with EtOAc (3 × 20 mL) and the combined fractions were washed with water (10 mL) then brine (10 mL) then dried ($Na_2SO_4$) and concentrated under reduced pressure to afford 1-(4-bromobenzyl)-N$^{\alpha}$-(*tert*-butoxycarbonyl)-N$^{\alpha}$-methyl-L-tryptophan (7.61 g, 83%) as a pale yellow solid. **$^1$H-NMR** (400 MHz; $CDCl_3$): δ 7.65 (d, *J* = 7.6 Hz, 1H), 7.51 (d, *J* = 8.4 Hz, 1H), 7.44-7.41 (m, 2H), 7.21 (dd, *J* = 6.1, 1.2 Hz, 2H), 6.95 (d, *J* = 8.4 Hz, 2H), 5.24 (s, 2H), 4.89-4.82 (m, 1H), 3.49-3.40 (m, 2H), 3.24-3.20 (m, 1H), 2.82 (s, 1.5H), 2.70 (s, 1.5H), 1.43 (s, 4.5H), 1.23 (s, 4.5H). **LCMS** (ESI): $[M-H]^-$, 486.1.

**N$^{\alpha}$-(((9H-Fluoren-9-yl)methoxy)carbonyl)-1-(4-bromobenzyl)-N$^{\alpha}$-methyl-L-tryptophan**

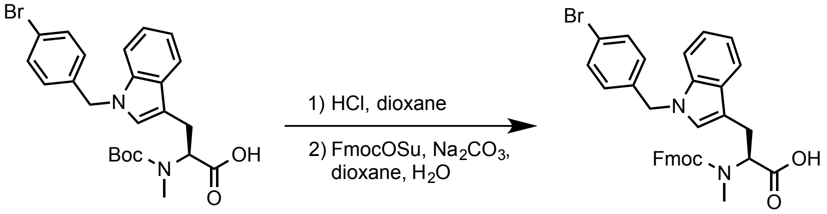


To a solution of 1-(4-bromobenzyl)-N$^{\alpha}$-(*tert*-butoxycarbonyl)-N$^{\alpha}$-methyl-L-tryptophan (2.50 g, 5.13 mmol) in dioxane (1 mL) was added HCl (5 mL of a 4 M solution in dioxane). The mixture was stirred for 2 h at room temperature then concentrated under reduced pressure, co-evaporated with dioxane several times then redissolved in a mixture of dioxane (12 mL) and water (12 mL). To this mixture was added $Na_2CO_3$ (1.25 g, 11.80 mmol) followed by FmocOSu (2.25 g, 6.67 mmol) and the mixture was stirred vigorously overnight. The mixture was concentrated to remove dioxane, acidified with citric acid (10 mL of a 10% aqueous solution) and extracted with EtOAc (3 × 10 mL) the combined organic fractions were washed with water, then brine, dried ($MgSO_4$), filtered, and concentrated under reduced pressure. The residue was subjected to flash chromatography (silica, gradient elution, 0-5% MeOH/DCM) to afford N$^{\alpha}$-(((9H-fluoren-9-yl)methoxy)carbonyl)-1-(4-bromobenzyl)-N$^{\alpha}$-methyl-L-tryptophan (2.83 g, 91%) as a pale yellow solid. **$^1$H-NMR** (400 MHz; $CDCl_3$, rotamers): δ 7.80-7.73 (m, 3H), 7.67-7.60 (m, 1H), 7.54 (t, *J* = 6.8 Hz, 1H), 7.49-7.39 (m, 5H), 7.36-7.31 (m, 2H), 7.22-7.12 (m, 4H), 6.97 (s, 1H), 6.91-6.86 (m, 2H), 5.18-5.15 (m, 2H), 5.05-5.01 (m, 0.5H), 4.87-4.82 (m, 0.5), 4.51-4.35 (m, 2H), 4.26-4.17 (m, 1.5H), 4.05-4.01 (m, 0.5), 3.56-3.50 (m, 1H), 3.41-3.34 (m, 1H), 2.86 (s, 3H). **LCMS** (ESI): $[2M-H]^-$, 1218.2.

**Allyl L-leucinate**

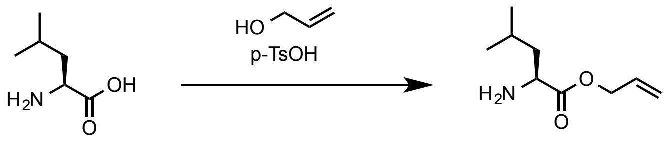


To a solution of L-leucine (2.50 g, 19.06 mmol) in allyl alcohol (25 mL) was added p-TsOH•$H_2O$ and the mixture was stirred at 90 °C (oil bath) for 16 h. The next day the solution was concentrated under reduced pressure to a solid residue, which was dissolved in DCM (25 mL) and washed with $NaHCO_3$ (10 mL of a saturated aqueous solution). The aqueous fraction was extracted with DCM (2 × 10 mL) and the combined organics were dried ($Na_2SO_4$), and filtered directly into HCl (4 equiv, ~ 19 mL of a 4 M solution in dioxane). The solution was concentrated under reduced pressure then precipitated with $Et_2O$. The mixture was briefly sonicated, left to settle for a few minutes, and the solution was decanted from the solid using a pipette. The product was dried under vacuum overnight, affording the title compound (3.07 g, 77%) as a pale yellow solid. **$^1$H-NMR** (400 MHz; DMSO-$d_6$): δ 8.61 (t, *J* = 0.4 Hz, 3H), 5.94 (ddt, *J* = 17.2, 10.6, 5.4 Hz, 1H), 5.39 (dq, *J* = 17.3, 1.6 Hz, 1H), 5.29 (dq, *J* = 10.5, 1.3 Hz, 1H), 4.70 (d, *J* = 5.4 Hz, 2H), 4.01-3.98 (m, 1H), 1.81-1.73 (m, 1H), 1.67 (t, *J* = 7.3 Hz, 2H), 0.91 (dd, *J* = 6.5, 1.2 Hz, 6H).

**Allyl N$^\alpha$-(((9H-fluoren-9-yl)methoxy)carbonyl)-1-(4-bromobenzyl)-N$^\alpha$-methyl-L-tryptophyl-L-leucinate**

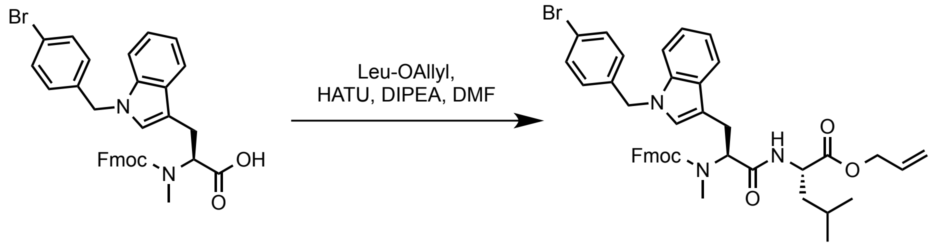


A 100 mL round bottom flask was charged with N$^\alpha$-(((9H-fluoren-9-yl)methoxy)carbonyl)-1-(4-bromobenzyl)-N$^\alpha$-methyl-L-tryptophan (6.50 g, 10.66 mmol), Leu-OAllyl (2.54 g, 12.26 mmol) and HATU (4.87 g, 12.87 mmol). The mixture was dissolved in DMF (50 mL) under argon and cooled to 0 °C, then DIPEA (5.57 mL, 32.00 mmol) was added and the mixture was stirred for 2 h, diluted with EtOAc (50 mL) and washed with water (4 × 20 mL). The combined aqueous phases were extracted again with EtOAc (10 mL) and the combined organics were washed with brine (10 mL) and dried ($Na_2SO_4$), filtered and concentrated to afford Allyl $N^{\alpha}$-(((9H-fluoren-9-yl)methoxy)carbonyl)-1-(4-bromobenzyl)-$N^{\alpha}$-methyl-L-tryptophyl-L-leucinate (7.92 g, 97%) a yellow solid. **$^1$H-NMR** (400 MHz; $CDCl_3$, rotamers): δ 7.80-7.79 (m, 2H), 7.73-7.63 (m, 2H), 7.55-7.34 (m, 5H), 7.18 (d, *J* = 0.1 Hz, 3H), 7.00 (s, 1H), 6.90-6.88 (m, 2H), 6.45-6.43 (m, 1H), 6.04-5.86 (m, 1H), 5.38-5.07 (m, 4H), 4.68-4.60 (m, 3H), 4.44-4.35 (m, 1H), 4.23-4.14 (m, 1H), 3.93-3.80 (m, 1H), 3.52-3.43 (m, 1H), 3.28-3.22 (m, 1H), 2.94-2.90 (m, 3H), 1.69-1.51 (m, 4H), 0.99-0.88 (m, 6H). **LCMS** (ESI): $[M+H]^+$, 764.6.

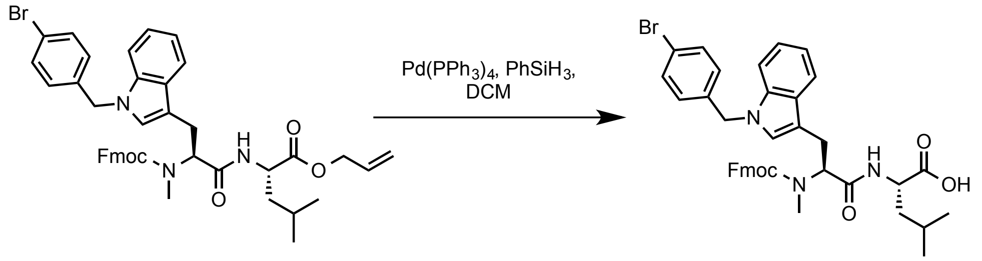


To a solution of the allyl ester (8.29 g, 10.87 mmol) in DCM (75 mL) at room temperature under argon was added $PhSiH_3$ (2.62 mL, 21.74 mmol) followed by $Pd(PPh_3)_4$ (628 mg, 0.54 mmol). The mixture was stirred for 2 h then concentrated onto silica and chromatographed (silica, 0-5%, MeOH/DCM) to afford the carboxylic acid (7.39 g, 94%) as a brown solid. **$^1$H-NMR** (400 MHz; $CDCl_3$, rotamers): δ 7.79 (dd, *J* = 7.2, 0.6 Hz, 2H), 7.66 (d, *J* = 1.4 Hz, 3H), 7.50-7.39 (m, 6H), 7.18-7.15 (m, 3H), 6.97 (d, *J* = 0.3 Hz, 1H), 6.87-6.85 (m, 2H), 5.18-5.06 (m, 3H), 4.60-4.54 (m, 1H), 4.39-4.31 (m, 1H), 4.21-4.14 (m, 1H), 3.54-3.40 (m, 2H), 3.25-3.18 (m, 1H), 2.94-2.84 (m, 3H), 1.72-1.50 (m, 3H), 0.94-0.87 (m, 6H). **LCMS** (ESI): $[M+H]^+$, 722.6.

**General procedures for peptide elongation on resin**

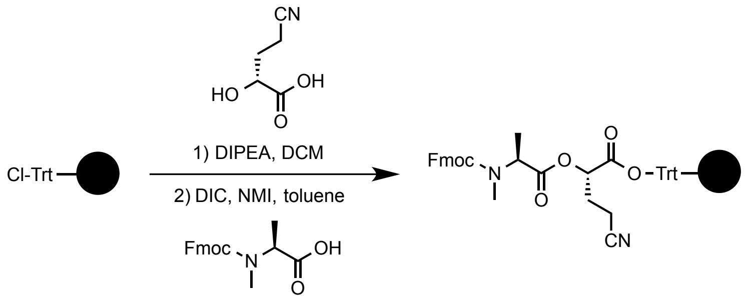


**Resin loading:**

A 50 mL Luer lock polypropylene syringe, fitted with a Teflon stopcock and internal polypropylene frit was charged with Cl-2-Cl-Trityl resin (2.5 g, Bachem, 1.6 meq/g, 4.07 mmol). The stopcock was closed, DCM (20 mL, anhydrous) was added and the syringe was fitted with a polypropylene plunger and agitated, end-over-end, for 1 h to swell the resin. The resin was filtered under suction and a solution of (2*R*)-4-cyano-2-hydroxybutanoic acid (789 mg, 6.11 mmol, 1.5 equiv) and DIPEA (2.11 mL, 12.23 mmol, 3 equiv) in DCM (20 mL, anhydrous) was added and the mixture was agitated, end-over-end, for 16 h. The resin was then filtered under suction and washed with DCM (2 × 20 mL × 1 min gentle shaking) then with toluene (anhydrous, 2 × 20 mL × 1 min gentle shaking). The resin was filtered under suction and a premixed solution of Fmoc-*N*-Me-Ala-OH (3.58 g, 11.0 mmol, 2.0 equiv), *N*,*N*-diisopropyl carbodiimide (1.72 mL, 11.0 mmol, 2 equiv) and *N*-methylimidazole (0.88 mL, 11.0 mmol, 2 equiv) in toluene (20 mL, anhydrous) was added and the mixture was agitated, end-over-end, for 1 h. The resin was drained and the coupling procedure was repeated once more. The resin was then washed using resin 'washing method A' described below. The resin was then dried under high vacuum overnight affording dried resin (3.59 g, 60%). Resin loading was also assessed by measuring absorbance at 290 nm of a solution prepared by treating a 1 mg sample of resin with 3 mL of a 20% 4-methylpiperidine solution in DMF for 10 min. The measured quantity of dibenzofulvene adduct was determined using a standardized calibration plot. Using this method, the measured loading was 0.96 mmol/g (60%).

**Resin washing method A:**
To the resin is added DMF (10 mL/g resin) and the resin is agitated by gently shaking for about 1 min. The resin is then drained under light vacuum and the procedure is repeated with *i*-PrOH (1 × 10 mL/g resin), then DMF (1 × 10 mL/g resin), then *i*-PrOH (1 × 10 mL), then DMF (1 × 10 mL/g resin), then DCM (3 × 10 mL/g resin).

**Resin washing method B:**
To the resin is added DMF (10 mL/g resin) and the resin is agitated by gently shaking for about 1 min. The resin is then drained under light vacuum and the procedure is repeated with *i*-PrOH (1 × 10 mL/g resin), then DMF (1 × 10 mL/g resin), then *i*-PrOH (1 × 10 mL/g resin), then DMF (3 × 10 mL/g resin).

**Resin washing method C:**
To the resin is added DMF (10 mL/g resin) and the resin is agitated by gently shaking for about 1 min. The resin is then drained under light vacuum and the procedure is repeated with *i*-PrOH (1 × 10 mL/g resin), then DMF (1 × 10 mL/g resin), then *i*-PrOH (1 × 10 mL/g resin), then DMF (1 × 10 mL/g resin), then toluene (3 × 10 mL/g resin).

**Fmoc removal:**
A solution of 4-methylpiperidine in DMF (20%, 10 mL/g resin) is added to the syringe containing the

DMF (or DCM)-swelled resin. The syringe is capped with a polypropylene plunger and the mixture was agitated, end-over-end, for 5 min. The resin is filtered under suction and the procedure repeated twice more. The resin is then drained under light vacuum and washed using ‘resin washing method B’.

**Resin coupling method A:**
To a solution of Fmoc-AA-OH (2 equiv) and HATU (2 equiv) in DMF (0.1 M) is added DIPEA (4 equiv) and the solution is quickly mixed to homogeneity and added to the syringe containing DMF-swelled resin. The syringe is capped with a polypropylene plunger and the mixture is agitated, end-over-end, for 1 h at room temperature, then drained under light vacuum and washed using ‘resin washing method B’. Chloranil staining assessed complete coupling: To ~0.5 mg of wet resin is added 20% acetaldehyde in DMF (100 µL) followed by 20 mg/mL chloranil in DMF (100 µL) and the beads are incubated for 2-5 min. Colorless beads indicated quantitative coupling for each step.

**Resin coupling method B:**
To a solution of Fmoc-AA-OH (2 equiv.) and EEDQ (2 equiv.) in toluene (0.35 M) and the solution is quickly mixed to homogeneity and added to the syringe containing toluene-washed resin. The syringe is capped with a polypropylene plunger and the mixture is agitated, end-over-end, for 2 h at room temperature, then drained under light vacuum and washed using ‘resin washing method B’. Chloranil staining assessed complete coupling: To ~0.5 mg of wet resin is added 20% acetaldehyde in DMF (100 µL) followed by 20 mg/mL chloranil in DMF (100 µL) and the beads are incubated for 2-5 min. Colorless beads indicated quantitative coupling for each step.

**Resin coupling method C:**
A solution of Fmoc-AA-OH (2 equiv.) and EEDQ (2 equiv.) in toluene (0.35 M) was sonicated to a thick gel. This gel was dissolved in DMF (final concentration 0.117 M). The solution was added to the syringe containing toluene-washed resin. The syringe is capped with a polypropylene plunger and the mixture is agitated, end-over-end, for 2 h at room temperature, then drained under light vacuum and washed using ‘resin washing method B’. Chloranil staining assessed complete coupling: To ~0.5 mg of wet resin is added 20% acetaldehyde in DMF (100 µL) followed by 20 mg/mL chloranil in DMF (100 µL) and the beads are incubated for 2-5 min. Colorless beads indicated quantitative coupling for each step.

**Synthesis of the linear peptide**

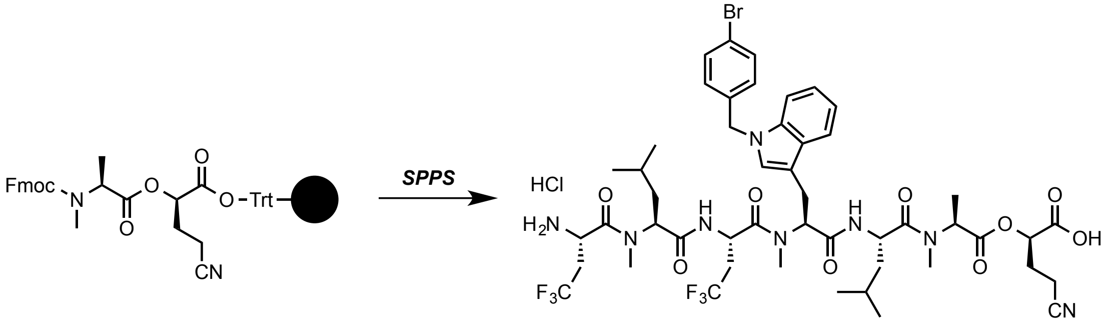

To dry Fmoc-N-Me-Ala-DGCN-loaded resin (3.00 g, 2.46 mmol) in the polypropylene syringe, fitted with a frit and stopcock, was added DCM (25 mL) and the syringe was capped with plunger and the resin was swelled by agitating, end-over-end, for 1 h. The resin was then drained and Fmoc group was removed using 'Fmoc removal' procedure. The resin was then washed using 'resin washing method C' then coupled with Fmoc-N-Me-Trp(4-BrBn)-Leu-OH using 'resin coupling method B', and washed with 'resin washing method B', then Fmoc was removed using 'Fmoc removal' procedure and resin washed with 'resin washing method B'. The peptide was elongated in a similar manner using:

1) Fmoc-Gly($CH_2CF_3$)-OH using 'resin coupling method C'
2) Fmoc-N-Leu-OH using 'resin coupling method A'
3) Fmoc-Gly($CH_2CF_3$)-OH using 'resin coupling method C'

After removal of the last N-terminal Fmoc group, the resin was washed using 'washing method A'.

**Cleavage from resin and isolation of heptadepsipeptide hydrochloride:**

To the syringe containing the washed heptadepsipeptide-loaded resin (assumed 2.46 mmol) obtained above was added HFIP (15 mL of a 20% solution in DCM) and the syringe was capped with a plunger and agitated, end-over-end, for 15 min (the resin turned red). Using a light vacuum, the solution was eluted into a 250 mL round bottom flask containing HCl (0.25 M solution in EtOAc, 40 mL ~4 equiv). The cleavage procedure was repeated once more and the solution was collected in the same flask of HCl. The remaining resin was rinsed into the HCl solution with DCM (2 × 10 mL) and the combined solution was concentrated under reduced pressure. The residue was dissolved in a minimal volume of EtOAc and $Et_2O$ was added and was accompanied by precipitation of a white solid. The mixture was sonicated for ~10 seconds and the precipitate left to settle for a few minutes before the liquid was carefully removed using a pipette. The residual solid was triturated with $Et_2O$ and dried under high vacuum to afford the heptadepsipeptide hydrochloride (930 mg, 33%) a white solid. The product was used in the next step without purification. **HRMS** (ESI): Calculated for $[M-H]^-$ $C_{53}H_{87}N_8O_9$, 1101.3878; Found, 1101.3868.

**Macrocyclization method A**

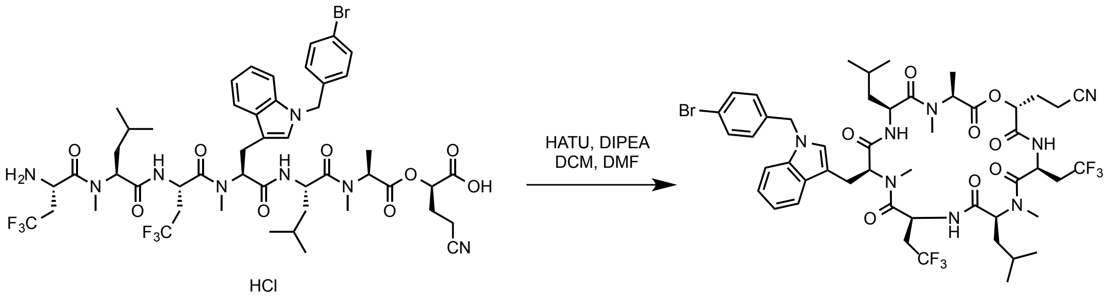


A solution of linear heptadepsipeptide (930 mg, 0.816 mmol) and DIPEA (427 µL, 2.45 mmol, 3 equiv.) in DCM (313 mL) was added, via a dropping funnel (at a rate of ~1 drop/sec), to a rapidly stirred solution of HATU (316 mg, 0.858 mmol), DMF (6 mL) and DCM (1.31 L). After complete addition, the funnel was rinsed into the reaction mixture with DCM (~10 mL) and the reaction was stirred for 18 h. An additional portion of HATU (158 mg, 429 mmol) was added and the reaction stirred for another 2 h. The reaction mixture was washed with HCl (500 mL of a 0.2 M aqueous solution), then with $NaHCO_3$ (500 mL of a 33% saturated aqueous solution). The DCM layer was then dried ($MgSO_4$), filtered and concentrated under reduced pressure. The residue purified by Flash chromatography (silica, step gradient elution, 5-50% acetone:hexane). Concentration of the appropriate fractions afforded **KZR-8445** (330 mg, 37%) as a white solid. **$^1$H-NMR** (400 MHz; acetone-$d_6$): δ 8.45 (d, *J* = 10.1 Hz, 1H), 8.25 (d, *J* = 9.6 Hz, 1H), 8.01 (d, *J* = 7.0 Hz, 1H), 7.71 (dd, *J* = 7.1, 0.9 Hz, 1H), 7.50-7.47 (m, 2H), 7.41 (t, *J* = 5.9 Hz, 1H), 7.36 (s, 1H), 7.18-7.09 (m, 4H), 5.45-5.34 (m, 2H), 5.26-5.06 (m, 3H), 4.76-4.71 (m, 1H), 4.37 (dd, *J* = 10.9, 3.6 Hz, 1H), 3.99-3.94 (m, 1H), 3.38-3.32 (m, 1H), 3.29 (s, 3H), 3.17-3.09 (m, 1H), 2.94 (s, 3H), 2.58 (s, 3H), 2.52-2.48 (m, 2H), 2.37-2.18 (m, 4H), 2.07 (dt, *J* = 4.4, 2.2 Hz, 2H), 1.96-1.82 (m, 2H), 1.57-1.45 (m, 8H), 1.04-0.94 (m, 10H), 0.90-0.85 (m, 2H), 0.49-0.42 (m, 1H). **LCMS** (ESI): Calculated for $C_{49}H_{62}BrF_6N_8O_8$ $[M+H]^+$, 1083.4; Found, 1083.2.

**KZR-9508** was synthesized using the procedures described above for the synthesis of **KZR-8445**. **$^1$H NMR** (600 MHz, DMSO-$d_6$) δ 8.86 (d, *J* = 6.7 Hz, 1H), 8.33 (d, *J* = 10.1 Hz, 1H), 8.05 (d, *J* = 9.6 Hz, 1H), 7.54 (d, *J* = 7.9 Hz, 1H), 7.41 (d, *J* = 6.9 Hz, 1H), 7.17 (s, 1H), 7.13 (t, *J* = 7.5 Hz, 1H), 7.02 (t, *J* = 7.4 Hz, 1H), 5.33–4.87 (m, 3H), 4.47–3.96 (m, 4H), 3.36 (s, 1H), 3.20–3.12 (m, 3H), 3.08–2.97 (m, 1H), 2.86–2.78 (m, 3H), 2.60 (dd, *J* = 12.9, 8.4 Hz, 1H), 2.43 (s, 3H), 2.35 (ddd, *J* = 17.5, 8.4, 3.5 Hz, 1H), 2.21 (d, *J* = 2.1 Hz, 1H), 2.03–1.60 (m, 4H), 1.53–1.21 (m, 14H), 0.94–0.84 (m, 12H), 0.30–0.15 (m, 1H). **LCMS** (ESI): Calculated for $C_{44}H_{61}F_6N_8O_8$+ $[M + H]^+$, 943.4511; Found 943.4492.
